# GATA2 controls lymphatic endothelial cell junctional integrity and lymphovenous valve morphogenesis through *miR-126*

**DOI:** 10.1101/660068

**Authors:** Md. Riaj Mahamud, Xin Geng, Yen-Chun Ho, Boksik Cha, Yuenhee Kim, Jing Ma, Lijuan Chen, Greggory Myers, Sally Camper, Debbie Mustacich, Marlys Witte, Dongwon Choi, Young-Kwon Hong, Hong Chen, Gaurav Varshney, James Douglas Engel, Shusheng Wang, Tae-Hoon Kim, Kim-Chew Lim, R. Sathish Srinivasan

## Abstract

Mutations in the transcription factor GATA2 cause lymphedema. GATA2 is necessary for the development of lymphatic valves (LVs) and lymphovenous valves (LVVs), and for the patterning of lymphatic vessels. Here, we report that GATA2 is not necessary for valvular endothelial cell (VEC) differentiation. Instead, GATA2 is required for VEC maintenance and morphogenesis. GATA2 is also necessary for the expression of cell junction molecules VE-Cadherin and Claudin5 in lymphatic vessels. We identified *miR-126* as a target of GATA2, and *miR-126^−/−^* embryos recapitulate the phenotypes of mice lacking GATA2. Primary human lymphatic endothelial cells (HLECs) lacking GATA2 (GATA2**^ΔHLEC^**) have altered expression of Claudin5 and VE-Cadherin, and blocking *miR-126* activity in HLECs phenocopies these changes in expression. Importantly, overexpression of *miR-126* in GATA2**^ΔHLEC^** significantly rescues the cell junction defects. Thus, our work defines a new mechanism of GATA2 and uncovers *miR-126* as a novel regulator of mammalian lymphatic vascular development.

**Non-standard abbreviations:** **LECs**, lymphatic endothelial cells; **LVs**, lymphatic valves; **LV-ECs**, lymphatic valve-forming endothelial cells; **LVVs**, lymphovenous valves; **LVV-ECs**, lymphovenous valve-forming endothelial cells; **HLEC**, primary human LECs; **OSS**, Oscillatory shear stress; **IHC**, immunohistochemistry.

## INTRODUCTION

The lymphatic vasculature is a hierarchically organized tissue that absorbs and returns extravasated plasma fluids and digested lipids to the blood circulation (Tammela and Alitalo 2010; Chen et al. 2014). This fluid, commonly known as lymph, is absorbed by lymphatic capillaries and transported via collecting lymphatic vessels. Lymphatic valves (LVs) within the lymphatic vessels regulate the unidirectional flow of lymph. Finally, lymph is returned to the blood circulation at the junction of jugular and subclavian veins through four lymphovenous valves (LVVs).

Mutations in multiple genes are associated with lymphedema, a debilitating disease characterized by the swelling of tissues, most obviously the limbs (Brouillard et al. 2014). Other lymphatic anomalies include chylous ascites (fluid in the peritoneal cavity), chylothorax (fluid around the lungs) or lymph reflex. These lymphatic vascular dysfunctions could arise from anatomical defects in the vessels or valves, though in most cases the precise cause is unclear.

Heterozygous mutations in the zinc finger transcription factor *GATA2* are associated with an array of hematopoietic disorders and lymphedema (Spinner et al. 2014). The overlapping phenotypes of this disease include immune deficiency, myelodysplasia (MDS), acute myeloid leukemia (AML), predisposition to mycobacterial infections and warts, hearing loss and lymphedema (Spinner et al. 2014; Crispino and Horwitz 2017). Emberger syndrome, caused by mutations in *GATA2*, is classified as deafness and primary lymphedema with MDS/AML (Emberger et al. 1979; Ostergaard et al. 2011; Kazenwadel et al. 2012). Approximately 11-30% of people with mutated *GATA2* develop lymphedema (Ostergaard et al. 2011; Kazenwadel et al. 2012; Spinner et al. 2014; Donadieu et al. 2018). Donadieu *et al*. noted that patients with *GATA2* mutations tend to develop lymphedema early, in the first decade of life. In summary, early-onset lymphedema with incomplete penetrance is associated with *GATA2*-heterozygous mutations. We need better insight into the molecular mechanisms of GATA2 activity to understand the causes of lymphedema in Emberger syndrome patients.

Mouse models have revealed that GATA2 is critical for the development of a variety of cell types including hematopoietic cells, neurons, pituitary glands, urinogenital system and the endothelium (Zhou et al. 1998; Zhou et al. 2000; Craven et al. 2004; Charles et al. 2006; Khandekar et al. 2007; Lim et al. 2012). *Gata2^−/−^* mice die at embryonic day (E)10 just as lymphatic endothelial cells (LECs) are starting to be specified. Conditional deletion of *Gata2* from all endothelial cells during mouse development results in severely edematous embryos with small blood-filled lymph sacs (Lim et al. 2012; Frye et al. 2018). Conditional deletion of *Gata2* in LECs results in mispatterned dermal lymphatic vessels, and a loss of lymphatic valves (LVs) (Kazenwadel et al. 2015; Frye et al. 2018). In addition, E12.5 or older embryos with a conditional deletion of *GATA2* in all endothelial cells or LECs lack LVVs (Kazenwadel et al. 2015; Geng et al. 2016; Frye et al. 2018). Thus, GATA2 is essential for proper development of the lymphatic vasculature.

*In vitro* experiments have revealed several molecular functions of GATA2. A stiff extra cellular matrix (ECM) triggers GATA2-dependent activation of *VEGFR2* expression in blood endothelial cells (Mammoto et al. 2009). In contrast, a soft-ECM enhances *GATA2* expression in primary human LECs (HLECs) and, in turn, induction of *VEGFR3* (Frye et al. 2018). This mechanistic relationship has been proposed to be critical for LEC migration from the cardinal vein and could explain the small lymph sacs in mice lacking *GATA2* in all endothelial cells.

Oscillatory shear stress (OSS), Wnt/β-catenin signaling and PROX1 are thought to be the most-upstream regulators of LV and LVV formation, all of which activate *GATA2* expression in HLECs (Kazenwadel et al. 2015; Sweet et al. 2015; Cha et al. 2016; Cha et al. 2018). OSS-induced GATA2 expression in HLECs is dependent on histone deacetylase 3 (HDAC3) (Janardhan et al. 2017). In turn, GATA2 is necessary for OSS-induced expression of FOXC2 and Connexin 37 (Kazenwadel et al. 2015; Sweet et al. 2015). Furthermore, GATA2 directly associates with the regulatory elements of PROX1 in HLECs, and GATA2 knockdown in HLECs downregulates the expression of PROX1 (Kazenwadel et al. 2015).

The current model built on these observations proposes that GATA2 regulates the differentiation of valvular endothelial cells from progenitors by upregulating PROX1, FOXC2 and Connexin 37 in those cells. However, whether this model is accurate *in vivo* remains unclear. Although LVV forming endothelial cells (LVV-ECs) differentiate at E12.0 with the upregulation of PROX1, FOXC2, Connexin 37 and GATA2 in those cells (Geng et al. 2016), whether GATA2 is necessary for LVV-EC differentiation is not known.

## RESULTS

### GATA2 is necessary for the proper architecture of newly differentiated LVV-ECs

Previous reports including ours have used pan-endothelial Cre lines for deleting *Gata2* (Kazenwadel et al. 2015; Geng et al. 2016; Frye et al. 2018). *Gata2* has also been deleted in the lymphatic vasculature in a mosaic manner using tamoxifen-inducible Cre lines (Kazenwadel et al. 2015; Frye et al. 2018). Here, we used *Lyve1-Cre* (Pham et al. 2010) to obtain uniform, constitutive deletion of *Gata2* (Charles et al. 2006) in the lymphatic vasculature. As anticipated *Lyve1-Cre;Gata2^f/f^* (*Gata2^LECKO^*) embryos recapitulated the previously reported lymphatic vascular phenotypes. Specifically, E16.5 *Gata2^LECKO^* embryos possessed blood-filled lymphatic vessels, which were dilated and had fewer branch points. The mutant embryos also lacked LVs and LVVs (**Figure 1** and data not shown).

**Figure 1:**
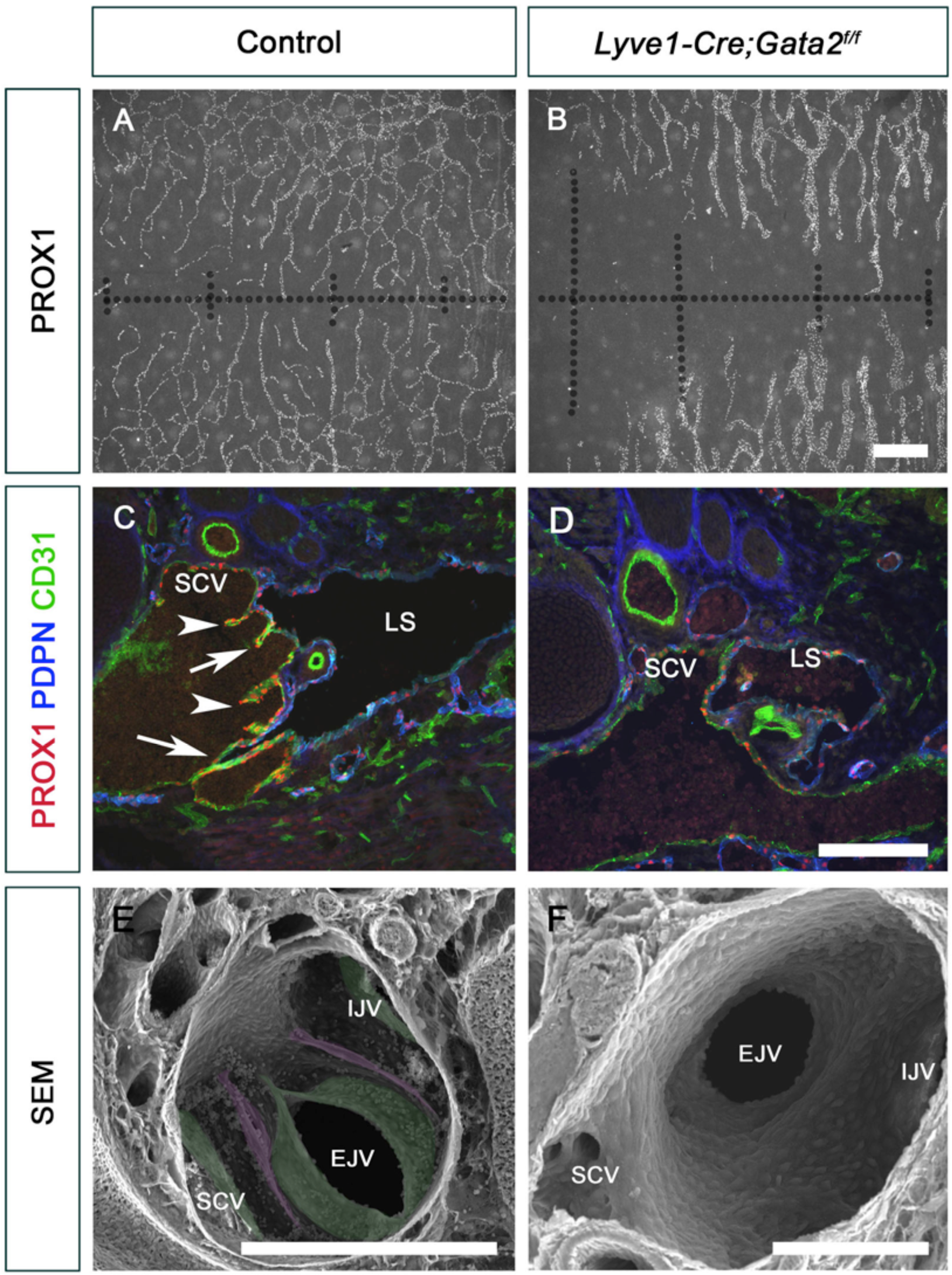
Lymphatic vessels are defective and LVVs are absent in *Lyve-Cre;Gata2^f/f^* embryos. E16.5 control and *Lyve-Cre;Gata2^f/f^* littermates were analyzed. (A, B) The lymphatic vessels in the dorsal skin of mutants were hypoplastic, dilated and had fewer branch points. (C, D) LVVs (arrows) and VVs (arrowheads) were seen in control, but not in mutants (D). (E, F) SEM confirmed that LVVs (magenta) and VVs (green) were present in control but not in mutant embryos. Abbreviations: IJV, internal jugular vein; EJV, external jugular vein; SCV, subclavian vein; LS, lymph sac. Measuring bar: (A-B) 500 µm; (C-D) 200 µm; (E) 300 µm; (F) 100 µm. Statistics: (A-B) n= 3 embryos; (C-D) n= 3 embryos and 6 LVVs per genotype; (E-F) n= 3 embryos and 5 LVV complexes per genotype.

To investigate a potential role for GATA2 in LVV-EC differentiation, we used numerous LVV-EC markers (PROX1^high^, FOXC2^high^, Connexin37 (GJA4), integrin-α5, integrin-α9) to analyze E12.0 embryos (**Figure 2A, B** and **Figure 2- Figure Supplement 1A-H**). LVV-EC numbers were not significantly different between control and *Gata2^LECKO^* embryos littermates at this stage (**Figure 2C**). We also analyzed E12.0 Tie2-Cre*;Gata2^f/f^* embryos in which *Gata2* is deleted from all endothelial cells at a much earlier time point (Kisanuki et al. 2001). E12.0 Tie2-Cre*;Gata2^f/f^* embryos had LVV-ECs (**Figure 2- Figure Supplement 2**), excluding the possibility that the LVV-ECs observed in *Gata2^LECKO^* embryos is due to inefficient deletion of *Gata2* by *Lyve1-Cre.* Thus, LVV-EC differentiation is normal in *Gata2^LECKO^* mutants.

**Figure 2:**
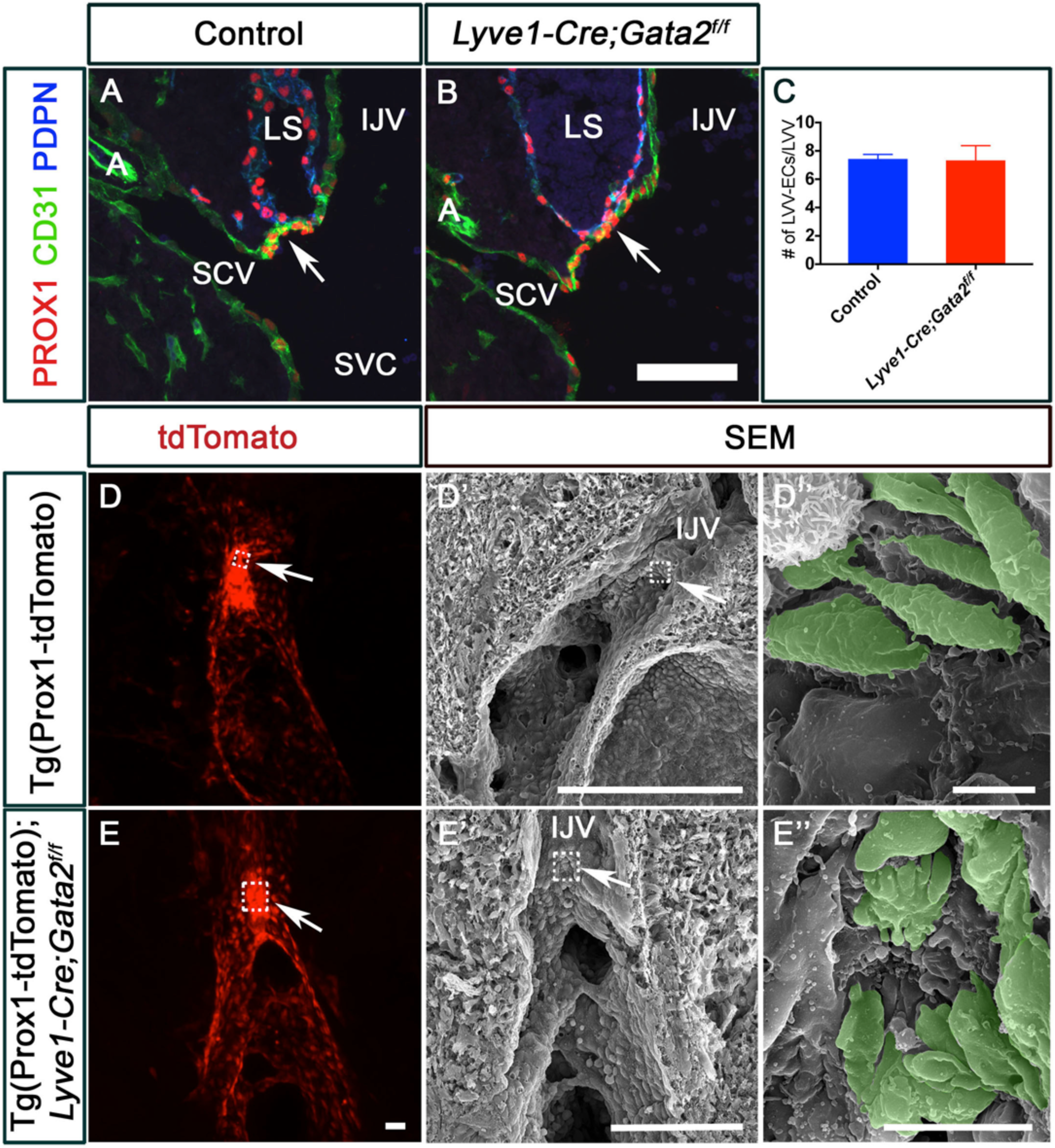
**GATA2 is required for the proper architecture of lymphovenous valve-forming endothelial cells (LVV-ECs).** (A-C) PROX1^high^ LVV-ECs were observed in both E12.0 control (A) and *Lyve1-Cre;Gata2^f/f^* (B) embryos. Blood cells were seen within the lymph sacs (LS) of mutant embryos. (C) No obvious difference in LVV-EC numbers was observed between the two genotypes. (D, E) E12.0 Tg(Prox1-tdTomato) (D) and Tg(Prox1-tdTomato); *Lyve1-Cre;Gata2^f/f^* (E) embryos were sagitally sectioned along the internal jugular vein (IJV). The fluorescent signal from the reporter revealed LVV-ECs in both control and mutant embryos. (D’, E’) The samples from D and E were analyzed using SEM, which revealed the LVV-ECs (pseudo-colored in green) with elongated morphology in control embryos (D’ and magnified figure of the boxed region in D’’). In contrast, the LVV-ECs of mutant embryos were dysplastic (E’ and E’’). Abbreviations: A, artery; IJV, internal jugular vein; SCV, subclavian vein; SVC, superior vena cava; LS, lymph sac. Measuring bar: (A, B) 100 µm; (D, E) 50 µm; (D’) 200 µm; (D”) 5 µm; (E’) 100 µm; (E”) 10 µm. Statistics: (A-E) n= 3 embryos and 6 LVV complexes per genotype per stage.

To examine LVV-ECs further, we used correlative fluorescent microscopy followed by scanning electron microscopy (SEM) to visualize the developing LVV-ECs at high resolution (Geng et al. 2016). First, we analyzed sagittal sections along the cardinal vein of E12.0 Tg(Prox1-tdTomato) embryos by confocal microscopy (Gong et al. 2003). We observed tdTomato^high^ LVV-ECs in both control and *Gata2^LECKO^* backgrounds (**Figure 2D, E**). SEM on these same samples revealed individual LVV-ECs that are elongated and aligned perpendicular to the direction of blood flow in control E12.0 embryos (**Figure 2D’’**, pseudo colored in green). The rest of the venous endothelium was quiescent with cobblestone-like appearance. In contrast, SEM revealed that the LVV-ECs in E12.0 Tg(Prox1-tdTomato); *Gata2^LECKO^* embryos are round and not aligned perpendicular to blood flow (**Figure 2- Figure Supplement 3)**. In addition, LVV-ECs also appear dysplastic in E12.0 Tg(Prox1-tdTomato); *Gata2^LECKO^* embryos (**Figure 2E’’**, pseudo colored in green, and **Figure 2- Figure Supplement 3)**. Based on these observations we conclude that GATA2 is not necessary for the differentiation of LVV-EC or for the upregulation of PROX1 or FOXC2 in those cells. However, GATA2 is necessary for the proper architecture of the newly formed LVV-ECs.

### GATA2 is necessary for the maintenance and morphogenesis of LVV-ECs

By E12.5, control embryos displayed LVV-ECs in the venous walls and two-well formed LVVs (**Figure 3A**, arrows). In contrast, E12.5 *Gata2^LECKO^* embryos had very few LVV-ECs in the venous walls and lacked clearly defined LVVs (**Figure 3B**, arrow). In addition, E12.5 Tg(Prox1-tdTomato) control embryos exhibited two tightly aggregated clusters of tdTomato^high^ LVV-ECs (**Figure 3C**, arrows), and SEM of one these clusters revealed elongated LVV-ECs that formed an opening in the middle (LVV) to permit lymph return to the blood circulation (**Figure 3C’’**, arrowhead). In contrast, Tg(Prox1-tdTomato);*Gata2^LECKO^* embryos lacked tdTomato^high^ LVV-ECs (**Figure 3D**), and SEM of the LVV-forming region revealed a smooth luminal surface, devoid of LVVs (**Figure 3D’’**).

**Figure 3:**
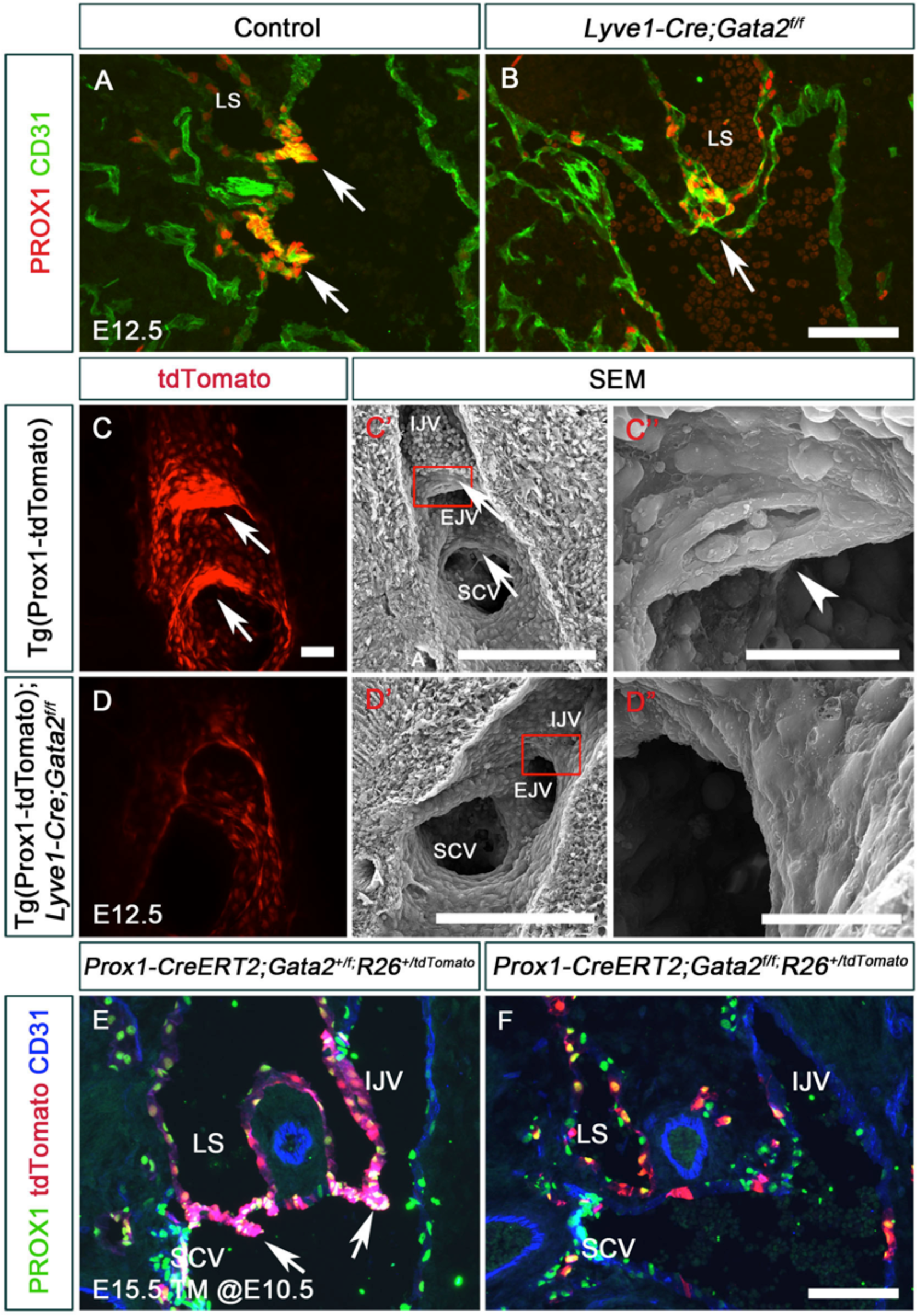
**LVV-ECs are lost from E12.5 embryos lacking GATA2.** (A, B) E12.5 wild type and *Lyve1-Cre;Gata2^f/f^* embryos were analyzed by immunohistochemistry on sections. LVV-ECs had invaginated into the vein in control embryos (A, arrows). In contrast, very few PROX1^+^ cells were observed in mutant embryos (B, arrow). (C, D) E12.5 Tg(Prox1-tdTomato) and Tg(Prox1-tdTomato); *Lyve1-Cre;Gata2^f/f^* embryos were sagitally sectioned along the internal jugular vein (IJV) and the fluorescent signal was analyzed using confocal microscopy. LVVs with strong tdTomato signal were seen in control (C, arrows), but not in mutant (D) embryos. (C’, C’’, D’, D’’) The samples from C and D were re-processed and analyzed by SEM. In controls, LVV-ECs (arrows) with elongated morphology were observed at the junction of IJV and external jugular vein (EJV) and at the junction of IJV and subclavian vein (SCV). An opening connecting the lymph and blood circulations was also seen (arrowhead in C’’). In contrast, endothelial cells at the junction of veins were indistinguishable from the rest of the venous endothelial cells in mutant embryos (D’, D’’). (E, F) Lineage tracing was performed using *Prox1-CreERT2*;*R26^+/tdTomato^* in *Gata2^+/f^* (E) or *Gata2^f/f^* (F) backgrounds. Tamoxifen was injected at E10.5 to label the PROX1^+^ LVV-ECs and LECs with tdTomato. Subsequently, the embryos were analyzed at E15.5. While LVV-ECs were well labeled in control embryos (E, arrows), LVVs were absent and very few labeled cells were observed in the veins of mutant embryos (F). Abbreviations: IJV, internal jugular vein; SCV, subclavian vein; SVC, superior vena cava; LS, lymph sac. Measuring bar: (A,B, E, F) 100 µm; (C, D) 50 µm; (C’) 200 µm; (D’) 500 µm; (C”, D”) 30 µm. Statistics: (A-B) n=6 embryos per genotype; (C-D) n= 3 embryos and 6 LVV complexes per genotype; (E, F) n=3 embryos per genotype.

Fluorescent reporter proteins such as tdTomato have a long half-life and can remain within cells for several days after the reporter gene is shut off (Muzumdar et al. 2007). However, tdTomato^high^ LVV-ECs in Tg(Prox1-tdTomato);*Gata2^LECKO^* embryos disappear within 12 hrs (between E12.0 to E12.5), suggesting that LVV-ECs were eliminated either by cell death or by detachment and removal via the bloodstream. To verify the loss of LVV-ECs, we performed lineage tracing. We generated *Prox1-CreERT2;R26^+/tdTomato^* embryos in a control (wild-type) and *Gata2^f/f^* background, treated pregnant dams with tamoxifen at E10.5, and evaluated embryos at E15.5. *R26^+/tdTomato^* allowed us to lineage trace the PROX1^+^ cells (LECs and LVV-ECs). Whereas entire LVVs were tdTomato^+^ in control embryos (**Figure 3E**, arrows), LVVs were absent in *Prox1-CreERT2;Gata2^f/f^;R26^+/tdTomato^* embryos (**Figure 3F**). Importantly, the LVV-forming area of *Prox1-CreERT2;Gata2^f/f^;R26^+/tdTomato^* embryos had very few labeled cells (**Figure 3F**), consistent with a loss of LVV-ECs in embryos lacking *Gata2*.

LVV-ECs in E12.0 or E12.5 control embryos did not express the proliferation marker phospho-histone 3 (PHH3) (data not shown), indicating that these cells do not proliferate. Therefore, the lack of LVV-ECs in *Gata2^LECKO^* embryos does not reflect impaired proliferation. We observed a few activated Casp3^+^ apoptotic cells within the lymph sacs of *Gata2^LECKO^* embryos, but LVV-ECs did not appear to be labeled by this marker for apoptosis (data not shown). Together these results indicate that GATA2 is not required for the differentiation of LVV-ECs or for their survival or proliferation. We however cannot exclude the possibility of cell death with the simultaneous detachment of LVV-ECs into the bloodstream.

In summary, GATA2 regulates the morphology of LVV-EC, maintains their presence in the valve-forming region, and regulates their morphogenesis into LVVs.

### GATA2 is not necessary for the upregulation of PROX1 and FOXC2 in VVs, LVs or AoVs

GATA2 is also expressed in LVs, venous valves (VVs) and aortic valves (AoVs) (Kazenwadel et al and data not shown) (Kazenwadel et al. 2015). Hence, we tested whether GATA2 is necessary for the differentiation of those valvular endothelial cells. VVs of the jugular vein exist close to LVVs at E16.5. VV-forming endothelial cells (VV-ECs) differentiate at this region at around E14.5. Control E15.5 embryos displayed LVVs (**Figure 4A**, arrows) and developing VVs invaginating into the veins (**Figure 4A**, arrowheads). In contrast, *Gata2^LECKO^* E15.5 embryos lacked LVVs, and PROX1^high^ VV-ECs were not invaginating into the veins (**Figure 4B**, arrowheads). Expression of FOXC2 is also unaffected in the VV-ECs of *Gata2^LECKO^* embryos (data not shown). Thus, GATA2 is not necessary for the differentiation of VV-ECs.

**Figure 4:**
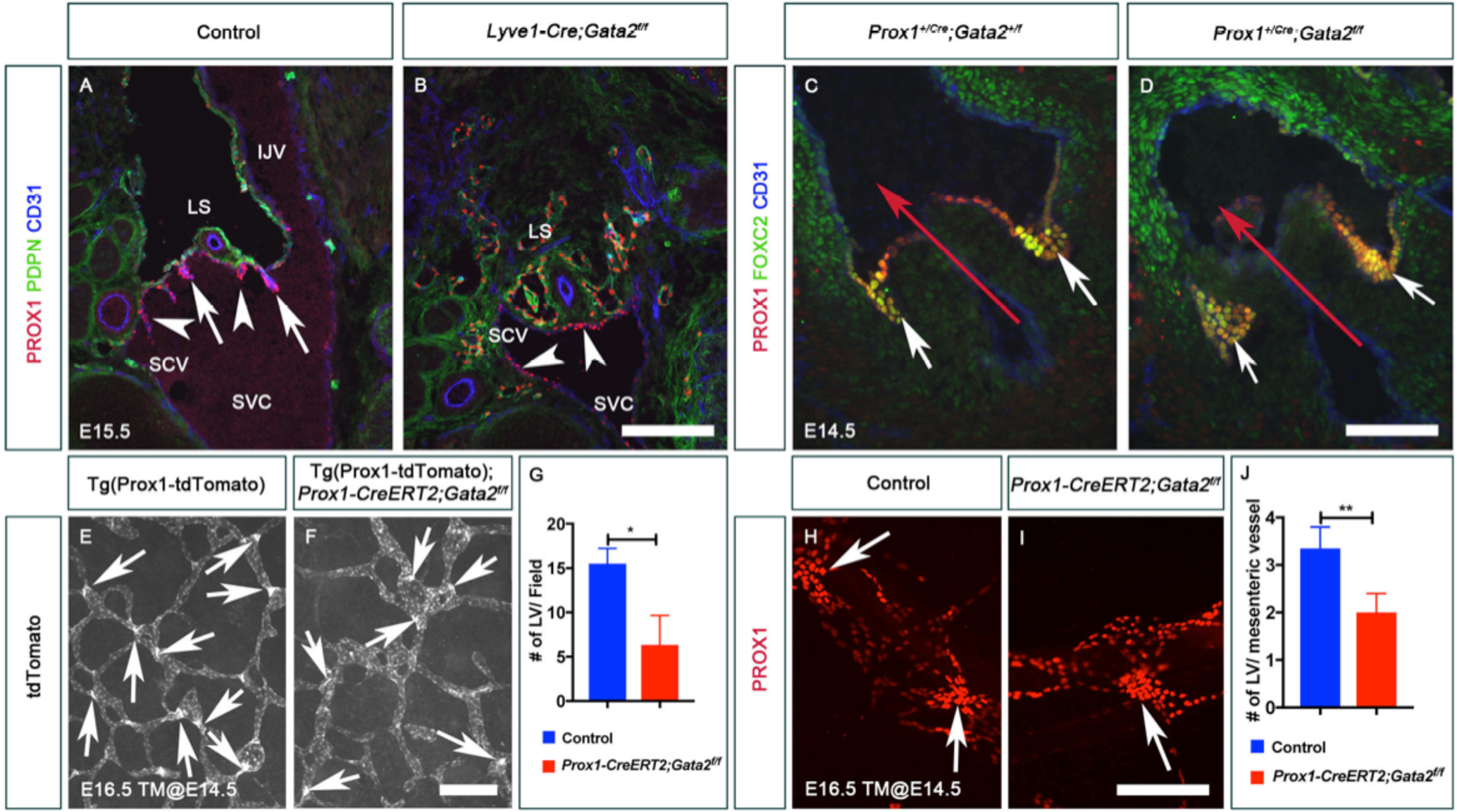
**GATA2 is not required for the differentiation of venous valve-, cardiac valve-, and lymphatic valve-forming endothelial cells.** (A) LVVs (arrows) and venous valves (VVs, arrowheads) were seen at the junction of internal jugular vein (IJV), subclavian vein (SCV) and superior vena cava (SVC) of E15.5 control embryos. (B) LVVs were absent in E15.5 *Lyve1-Cre;Gata2^f/f^* embryos. While VV-ECs were present in the mutants (arrowheads), they did not invaginate into the veins and did not have the proper morphology of VVs. (C, D) PROX1^+^ FOXC2^+^ endothelial cells were observed on the downstream side of cardiac valves in both control (C) and *Prox1^+/Cre^;Gata2^f/f^* (D) embryos, which lack GATA2 in all PROX1-expressing cells. The red arrow indicates the direction of blood flow. (E-J) Lymphatic vessels in the dorsal skin (E-G) and mesentery (H-J) of E16.5 control and *Prox1-CreERT2;Gata2^f/f^* embryos, which were exposed to tamoxifen at E14.5, were analyzed. Lymphatic valve-forming endothelial cells (LV-ECs) were seen in both control and mutant embryos (arrows). However, LV clusters are reduced in mutants (G, J). Abbreviations: IJV, internal jugular vein; SCV, subclavian vein; SVC, superior vena cava; LS, lymph sac. Measuring bar: (A, B, H, I) 200 µm; (D) 100 µm; (E, F) 250 µm. Statistics: (A-B) n= 3 embryos and 6 LVV complexes per genotype; (C-D) n= 3 embryos per genotype; (E-F, H-I) n= 3 embryos per genotype. (**) p<0.01; (*) p<0.05.

We deleted *Gata2* using *Prox1^+/Cre^* to remove GATA2 from the PROX1^+^ aortic valve endothelial cells (AoV-ECs) (Srinivasan et al. 2010). PROX1 and *Prox1^+/Cre^* are expressed in AoV-ECs as early as E12.5 (data not shown). As shown in **Figure 4C, D**, expression of PROX1 and FOXC2 was unaffected in the AoV-ECs of E14.5 *Prox1^+/Cre^;Gata2^f/f^* embryos. We also did not observe any obvious differences in the expression of PROX1 and FOXC2 in E16.5 *Prox1^+/Cre^;Gata2^f/f^* embryos (data not shown). Thus, GATA2 is not required to upregulate PROX1 and FOXC2 in AoV-ECs or to maintain these cells.

We noticed that in contrast to *Gata2^LECKO^* embryos *Prox1-CreERT2;Gata2^f/f^* embryos do not develop blood-filled lymphatic vessels even though they lack LVVs (**Figure 3F**). Blood-filled lymphatic vessels could affect LV development (Sweet et al. 2015). Therefore, we analyzed *Prox1-CreERT2;Gata2^f/f^* embryos to investigate the role of GATA2 in LV development. We exposed pregnant dams carrying *Prox1-CreERT2;Gata2^f/f^* embryos to tamoxifen at E14.5 and harvested the embryos at E16.5. Analysis of the skin and mesenteric lymphatic vessels revealed that LV-EC clusters were present in both control and mutant embryos, although those numbers were reduced in the mutants (**Figure 4E-J)**. In contrast, E18.5 *Prox1-CreERT2;Gata2^f/f^* embryos that were exposed to tamoxifen at E14.5 completely lacked LVs (data not shown). Hence, these results suggest that GATA2 is not necessary for the differentiation of LV-ECs, but it is necessary to maintain those cells.

Together, these results indicate that GATA2 is not necessary for the differentiation of LVV-ECs, VV-ECs, LV-ECs or AoV-ECs or for the upregulation of PROX1 and FOXC2 in those cells. However, GATA2 is necessary to maintain vascular valve endothelial cells (LVV-ECs and VV-ECs) and promote their morphogenesis.

### *EGFL7* and *ANGPT2* are regulated by GATA2 in HLECs

To identify potential genes regulated by GATA2 *in vivo*, we examined GATA2-dependent gene expression in HLECs. We treated HLECs with lentiviral particles expressing shGFP or shGATA2, harvested them 72 hrs later and performed RNA-seq. We performed this experiment in triplicate, and Principle Component Analysis confirmed the consistency in gene expression changes among the triplicates (**Figure 5A**). We pursued genes whose expression is significantly (p<0.05) different between shGFP- and shGATA2-treated HLECs, with Log_2_ fold change (FC) >0.5 or Log_2_ FC<-0.5. According to these criteria, 1009 genes were significantly downregulated and 617 genes were significantly upregulated upon depletion of *GATA2* in HLECs (**Figure 5B** and **Supplementary File 1**). *GATA2* was dramatically downregulated (Log_2_ FC = −2.08) in shGATA2-treated HLECs, as expected (**Supplementary File 1**). We did not observe significant changes in the expression of *PROX1, FOXC2 or FLT4*. However, a number of other genes that regulate vascular development were differentially expressed in shGATA2-treated HLECs (**Figure 5C**).

**Figure 5:**
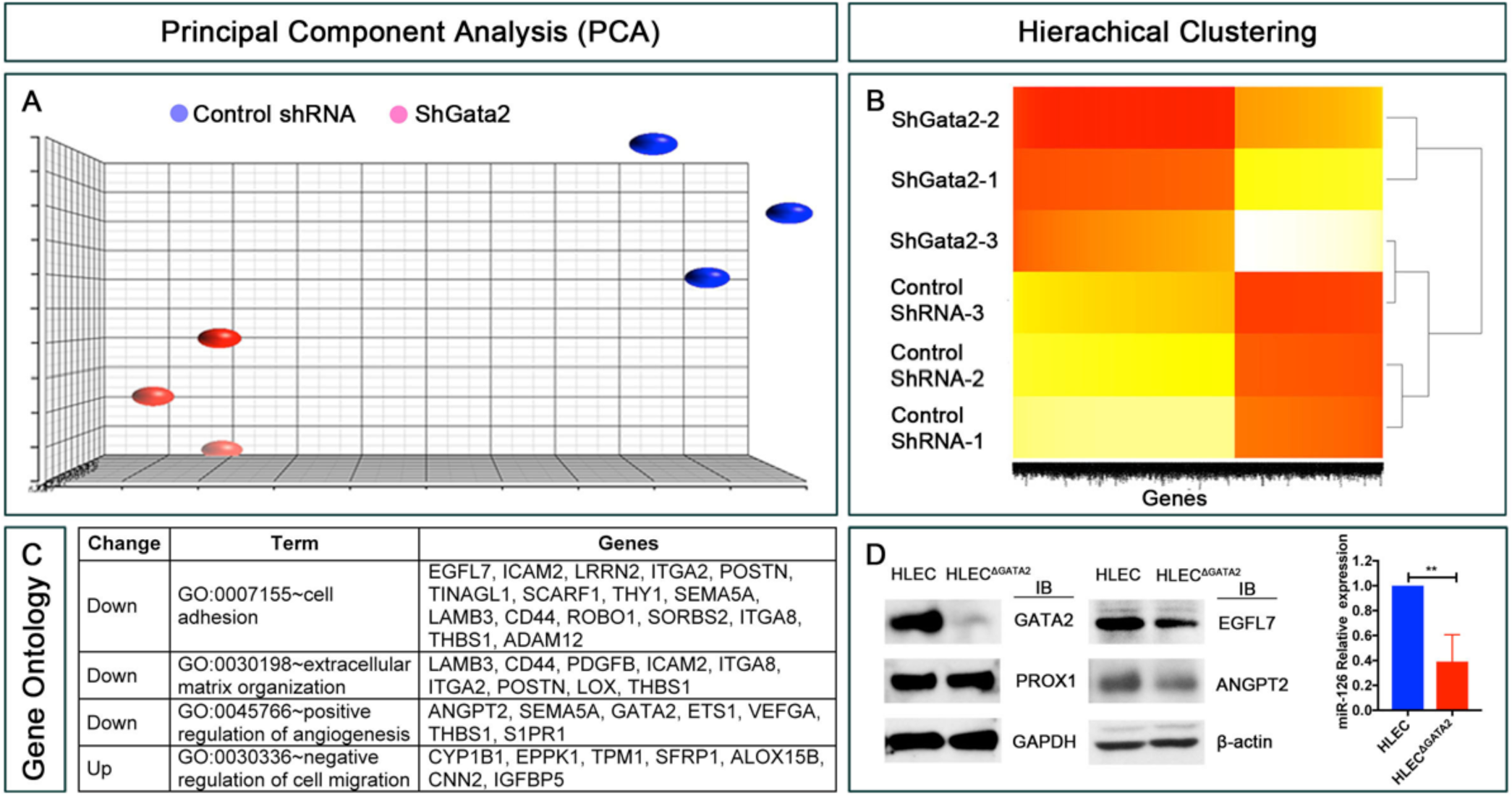
**RNA-seq identifies the targets of GATA2 in primary human LECs.** (A) Principal component analysis (PCA) was performed on RNA-seq data from control shRNA and shGATA2 infected primary human lymphatic endothelial cells (HLECs). A high level of similarity was observed within the groups as indicated by their proximity to each other. (B) Hierarchical clustering shows that approximately 1000 genes were consistently downregulated and 600 genes were upregulated in shGATA2 treated HLECs. (C) GO revealed a list of genes that are likely relevant to the phenotypes observed mice lacking GATA2. (D) GATA2 was knocked out from a second HLEC line using CRISPR/Cas9. Western blot revealed the lack of GATA2 and the downregulation of EGFL7 and ANGPT2 in the knock out cells (HLEC^ΔGATA2^). In contrast, no obvious differences were observed in the expression of PROX1. Additionally, qRT-PCR revealed the downregulation of miR-126. Statistics: (A) n=3 independent experiments per shRNA; (D) n= 3 independent experiments (antibiotic selection, Western blot and qRT-PCR). (**) p<0.01

Among the top 50 most down regulated genes only *GATA2*, *Angiopoietin 2* (*ANGPT2*) and *EGF-like domain-containing protein 7 (EGFL7*) are reported to be necessary for embryonic survival and vascular development (http://www.informatics.jax.org). *EGFL7* (NM_201446) is a secreted protein that was first reported as a regulator of vascular lumen formation (Parker et al. 2004). EGFL7 also regulates blood endothelial cell migration, contractility and adhesion (Charpentier et al. 2013). *EGFL7* is also the host gene for miR-126, which is the first endothelial cell-specific microRNA to be reported (Lagos-Quintana et al. 2002). Interestingly, EGFL7 is not required for the survival of mice as long as miR-126 is intact (Kuhnert et al. 2008). In contrast, most *miR-126^−/−^* embryos die in utero with severe edema (Kuhnert et al. 2008; Wang et al. 2008). Importantly, GATA2 was recently reported to regulate EGFL7 and miR-126 in blood vascular endothelial cells (Hartmann et al. 2016). However, the lymphatic vasculature specific roles of miR-126 remain unknown.

ANGPT2 is a secreted molecule and a ligand of TIE2 (also known as TEK). In blood endothelial cells, ANGPT2 is an antagonist of TIE2. The TIE2/ANGPT2 interaction in blood endothelial cells results in VE-PTP-mediated downregulation of VE-Cadherin (Souma et al. 2018). In contrast, in LECs ANGPT2 is an agonist of TIE2 due to the absence of VE-PTP. Deletion of *Angpt2* results in a strain-specific postnatal lethality in mice due to severe chylothorax (Gale et al. 2002; Dellinger et al. 2008). *Angpt2^−/−^* mice lack LVs and have defective cell junctions. Whether ANGPT2 is necessary for LVV development is unknown.

To validate whether ANGPT2 and EGFL7 are GATA2 targets, we used CRISPR/Cas9 to generate *GATA2* knockout in a distinct HLEC cell line (HLEC-2). Western blotting and DNA sequencing confirmed the deletion of *GATA2* in GATA2^ΔHLEC^ (**Figure 5D** and **Figure 5- Figure Supplement 1**). Importantly, the protein levels of both EGFL7 and ANGPT2 were reduced in GATA2^ΔHLEC^ cells compared to controls, whereas PROX1 was unaffected (**Figure 5D**). In addition, after extracting miRNA’s from the cells we determined by qRT-PCR that miR-126 is significantly downregulated in GATA2^ΔHLEC^ cells compared to controls (**Figure 5D**).

In summary, *EGFL7*, miR-126 and *ANGPT2* are consistently downregulated by the knockdown or knockout of GATA2 in HLECs.

### *MiR-126* is a physiologically important target of GATA2 in the lymphatic vasculature

To investigate the physiological relevance of these candidate GATA2 target genes, we compared their expression in the LVV-ECs of E12.0 control and *Gata2^LECKO^* embryos. ANGPT2 was not expressed in LVV-ECs at E12.0 although it appears at E14.5 (**Figure 6**). *Angpt2^−/−^* embryos lacked LVs and had defective lymphatic vessel patterning as reported previously (data not shown) (Gale et al. 2002; Dellinger et al. 2008). However, *Angpt2^−/−^* embryos retained normal looking LVVs and VVs (**Figure 6**). These observations suggested that GATA2-dependent regulation of *Angpt2* is not involved in LVV and VV development. Hence, we focused our attention on miR-126 for the rest of this work.

**Figure 6:**
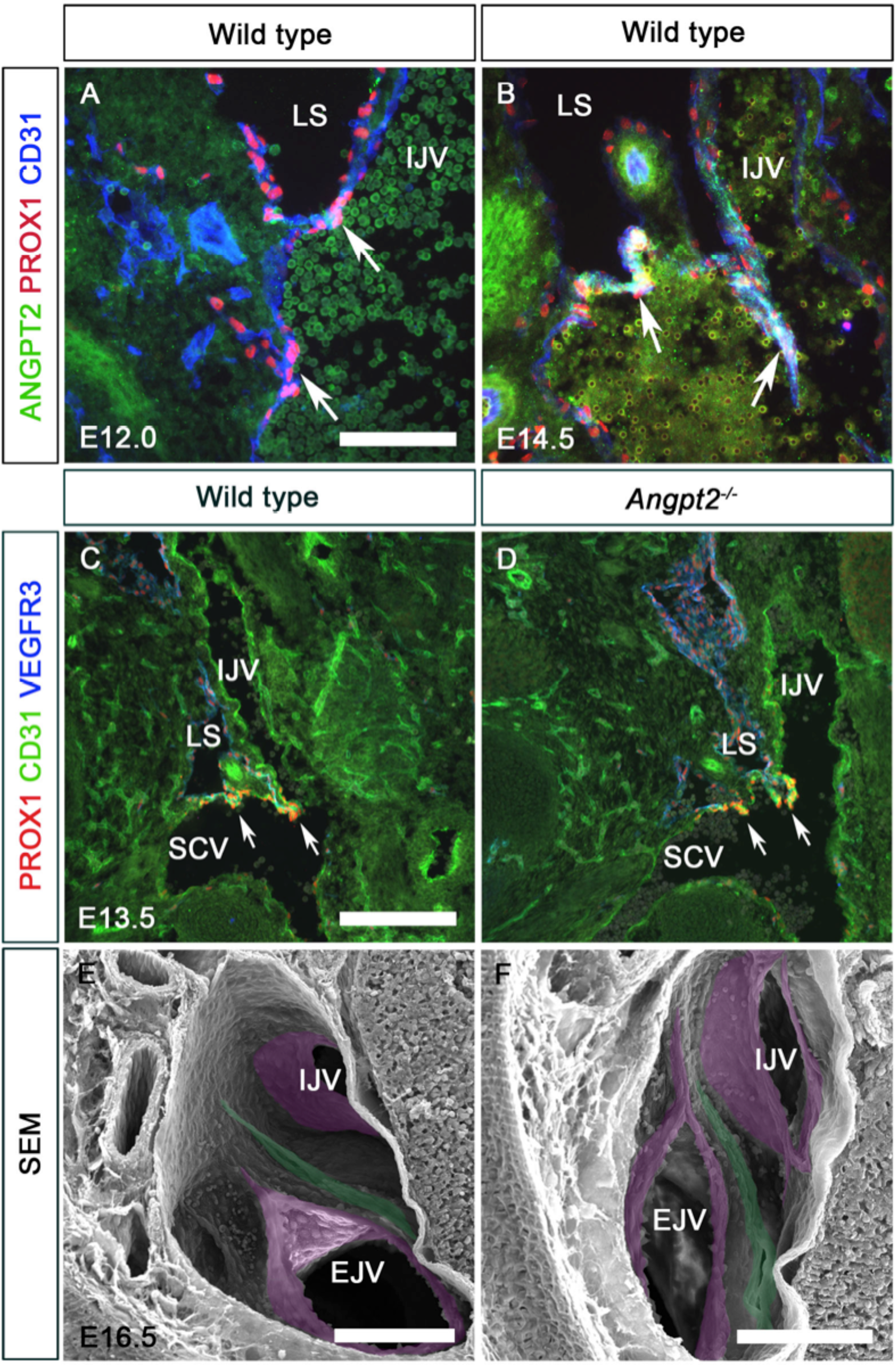
**Angiopoietin2 does not regulate LVV development.** (A, B) ANGPT2 was not expressed in E12.0 LVV-ECs (A, arrows). However, it was expressed in LVV-ECs at E14.5 (B, arrows). (C, D) LVVs developed normally in E13.5 *Angpt2^−/−^* embryos (arrows). (E, F) LVVs (green) and VVs (magenta) developed normally in E16.5 *Angpt2^−/−^* embryos. Abbreviations: IJV, internal jugular vein; EJV, external jugular vein; LS, lymph sac. Measuring bar: (A-B) 100 µm; (C-D) 200 µm; (E-F) 100 µm. Statistics:(A-F) n= 3 embryos and 6 LVV complexes per genotype.

*In situ* hybridization revealed high expression of *miR-126* in LVV-ECs of E12.0 control embryos but not *Gata2^LECKO^* embryos (**Figure 7A, B**, arrows). Similarly, E18.5 control embryos expressed EGFL7 in the LECs of mesenteric lymphatic vessels and in LVs (**Figure 7C**, arrow), whereas EGFL7 expression was dramatically reduced in the LECs of E18.5 *Prox1-CreERT2*;*Gata2^f/f^* embryos in which *Gata2* deletion was induced by tamoxifen injection at E14.5 (**Figure 7D**). As mentioned previously, the mutants lacked LVs. Thus, GATA2 is required for *EGFL7/miR-126* expression in the developing lymphatic vasculature.

**Figure 7:**
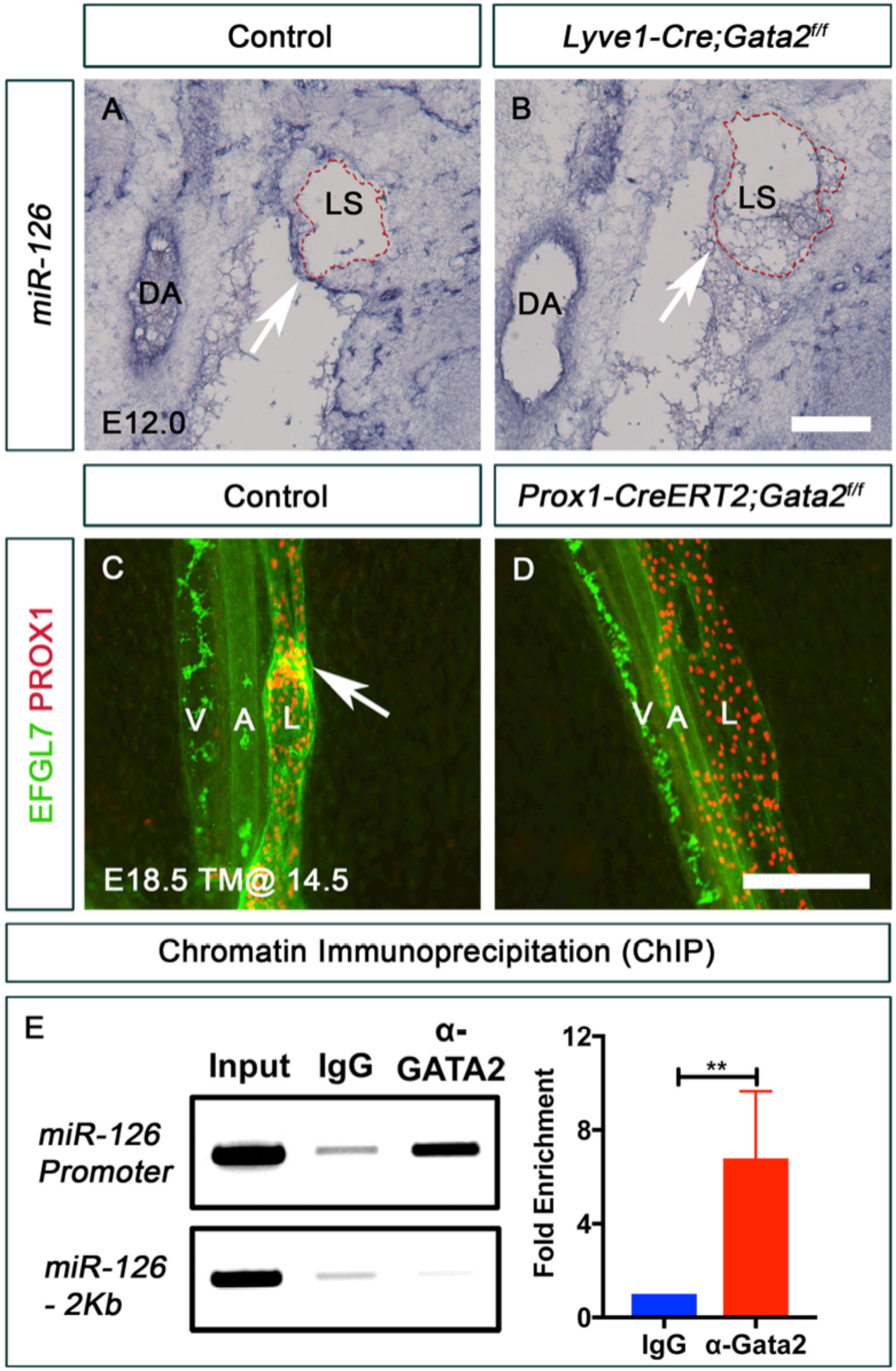
*EGFL7/miR-126* is a target of GATA2. (A) miR-126 was expressed in the dorsal aorta (DA) and in the LVV-ECs (arrow) of control embryos. (B) Expression of miR-126 was downregulated in the LVV-ECs (arrows) of E12.0 *Lyve1-Cre;Gata2^f/f^* embryos. However, no obvious difference in miR-126 expression was observed in the dorsal aorta (DA) of mutants. (C) EGFL7 was expressed in the mesenteric arteries (A), veins (V) and lymphatic vessels (L) of E18.5 control embryos. Strongest expression of EGFL7 was observed in LVs (arrow). (D) Expression of EGFL7 was dramatically downregulated in the mesenteric lymphatic vessels of mice lacking GATA2 in LECs. Also, notice the absence of LVs in the mutant. (E) Chromatin immunoprecipitation (ChIP) revealed that GATA2 strongly associates with the promoter element of the *EGFL7/miR-126* locus. The top lane of the gel picture was PCR performed using primers flanking the GATA2 binding site. The lower lane was PCR performed using primers for a non-specific site. The graph compares qPCR signals generated by primers flanking the GATA2 binding site. Measuring bar: (A, B) 250 µm; (C, D) 200 µm. Statistics: (A, B) n= 3 embryos and 6 LVV complexes per genotype; (C-D) n=3 embryos per genotype; (E) n=4. (**) p<0.01

GATA2 associates with the promoter of *EGFL7/miR-126* in human umbilical vein endothelial cells (HUVECs) (Hartmann et al. 2016). We performed chromatin immunoprecipitation (ChIP) using an anti-GATA2 antibody and determined that GATA2 associates with this promoter region in HLECs as well (**Figure 7E**). These results suggest that *EGFL7/miR-126* is a direct target of GATA2 in the lymphatic vasculature.

As mentioned previously *Egfl7^−/−^* mice that retain miR-126 are phenotypically normal (Kuhnert et al. 2008). Therefore, we analyzed *miR-126^−/−^* embryos, which display severe edema (Wang et al. 2008). LVVs and VVs were absent in the jugulo-subclavian vein junction of E16.5 *miR-126^−/−^* embryos compared to wild type (**Figure 8A, B**). LVV-ECs were present in E12.0 *miR-126^−/−^* embryos (**Figure 8C, D**), indicating that *miR-126* is not necessary for the differentiation of LVV-ECs, but for their maintenance. *MiR-126^−/−^* embryos had dilated mesenteric lymphatic vessels that lack LVs (**Figure 8E, F**), and the lymphatic vessels in the dorsal skin are hypoplastic (**Figure 8G, H**).

**Figure 8:**
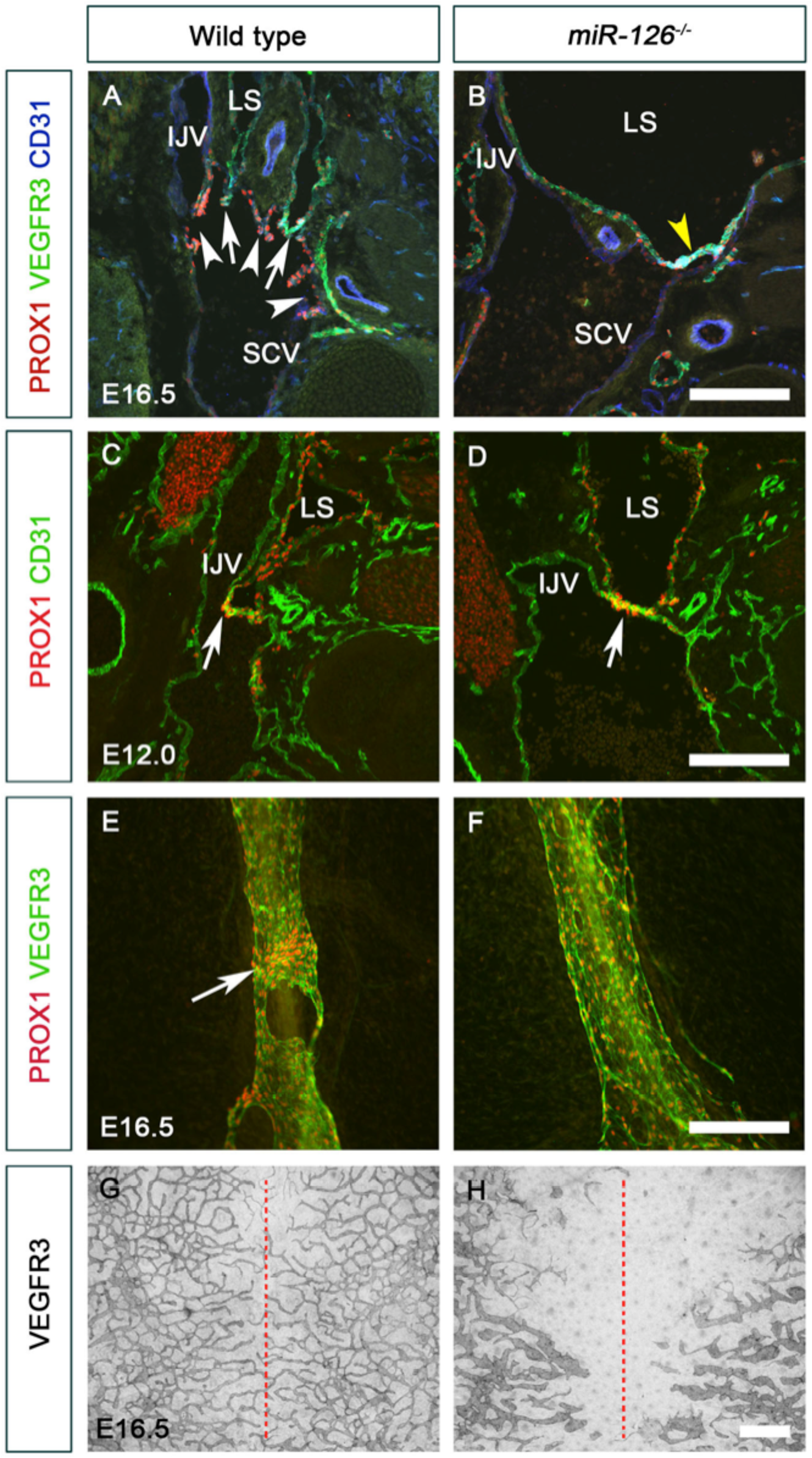
*miR-126^−/−^* embryos are phenotypically similar to mice lacking *Gata2* in LECs. (A, B) LVVs (arrows) and VVs (arrowheads) were seen at the junction of internal jugular vein (IJV) and subclavian vein (SCV) of E16.5 control (A) but not *miR-126^−/−^* (B) embryos. A few PROX1^+^ cells were nevertheless seen at the interface of vein and lymph sacs (B, yellow arrowhead). (C, D) LVV-ECs were observed in both E12.0 control and *miR-126^−/−^* littermates (arrows) indicating that miR-126 is not necessary for the differentiation of LVV-ECs. (E, F) LVs were observed in the mesenteric lymphatic vessels of E16.5 control embryos (E, arrow). PROX1 expression was higher in the LVs compared to LECs. (F) LVs were absent and PROX1 expression was homogeneous in the LECs of *miR-126^−/−^* littermates. (G, H) Lymphatic vessels in the dorsal skin had migrated from the lateral edges until the midline (red dotted lines) in control (G) but not the *miR-126^−/−^* (H) embryos. In addition, the lymphatic vessels of the mutants were dilated with fewer branch points. Abbreviations: IJV, internal jugular vein; SCV, subclavian vein; LS, lymph sac. Measuring bar: (A-F) 200 µm; (G, H) 1000 µm. Statistics: (A-D) n= 3 embryos and 6 LVV complexes per genotype; (E-H) n= 3 embryos per genotype.

Overall, these observations demonstrate that GATA2 regulates miR-126 both in vitro and in vivo. Furthermore, a significant level of phenotypic similarity between *Gata2^LECKO^* and *miR-126^−/−^* embryos, including defects in the maintenance of LVVs and lymphatic vessel patterning indicate that miR-126 is likely a physiologically relevant target of GATA2 in the lymphatic vasculature.

### GATA2 regulates lymphatic endothelial cell junctions via *miR-126*

The Ras MAPK pathway inhibitor *Sprouty related, EVH1 domain-containing protein 1 (Spred1)* is a target of miR-126 in blood endothelial cells (Fish et al. 2008; Wang et al. 2008), and deletion of *Spred1* rescues the blood vascular defects of *miR-126^−/−^* embryos (Wang et al. 2008; Zhou et al. 2016). To investigate a potential role for *Spred1* in the lymphatic vasculature, we examined *Spred1* knockout embryos. *miR-126^−/−^* embryos displayed edema as anticipated **(Figure 9A, B)**. *Spred1^−/−^* embryos did not have edema or any obvious defects in LVVs or lymphatic vessels **(Figure 9C-H)**. Importantly, deletion of *Spred1* did not rescue the lymphatic vascular defects of *miR-126^−/−^* embryos **(Figure 9C-H)**. Thus, miR-126 regulates lymphatic vascular development independently of *Spred1*.

**Figure 9:**
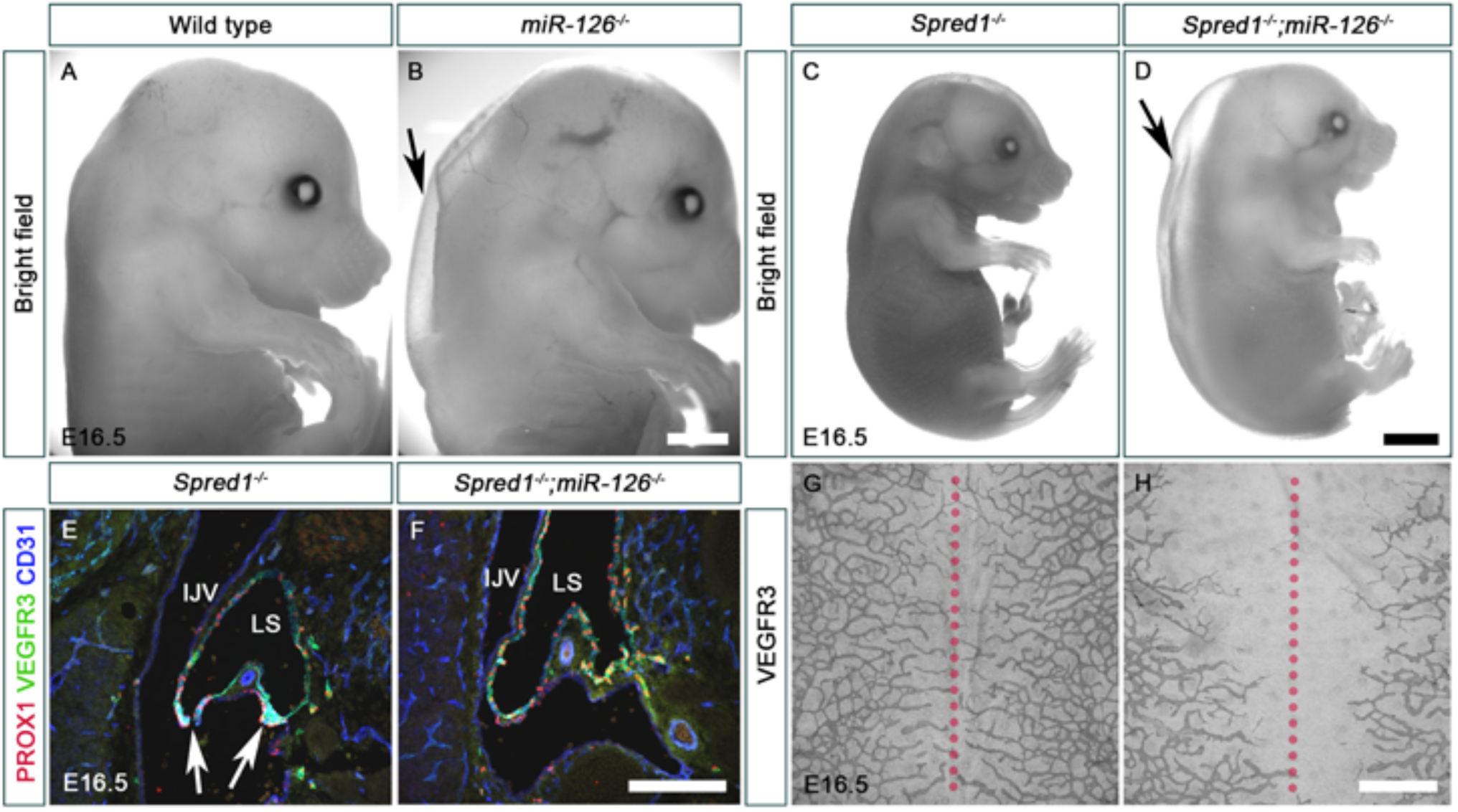
Deletion of Spred1 does not rescue the lymphatic vascular phenotypes of *miR-126^−/−^* embryos. (A-D) Severe peripheral edema (arrows) was observed in E16.5 *miR-126^−/−^* (B) and *miR-126^−/−^*; *Spred1^−/−^* (D) embryos. (C) E16.5 *Spred1^−/−^* embryos do not have obvious edema. (E-H) E16.5 *Spred1^−/−^* embryos had LVVs (E, arrows) and had normal lymphatic patterning (G). Deletion of *Spred1* does not rescue the absence of LVVs (F) or lymphatic vessel patterning defect (H) of *miR-126^−/−^* embryos. The dotted lines indicate the midline of dorsal skin. Abbreviations: IJV, internal jugular vein; LS, lymph sac. Measuring bar: (A, B) 5000 µm; (C, D) 2500 µm; (E, F) 200 µm; (G, H) 1000 µm. Statistics: (A-D, G-H) n= 3 embryos per genotype; (E, F) n=3 embryos, 6 LVV complexes per genotype.

To identify the relevant targets of miR-126 we performed RNA-seq in HLECs expressing an “miR-126 sponge” (Gentner et al. 2009; Lechman et al. 2012) to sequester *miR-126* from its endogenous targets (**Supplementary File 1**). Using the same criteria described above we identified 1058 genes that were upregulated and 873 genes that were downregulated by miR-126 sponge. SPRED1 and PIK3R2, which is another reported target of miR-126 in blood endothelial cells, were not in the list (Fish et al. 2008). *PROX1, FOXC2, FLT4* and *GATA2* were also not found in this list. By comparing these genes with the GATA2 regulated genes we identified 125 shared downregulated genes and 72 shared upregulated genes (data not shown). DAVID gene annotation was used to classify the shared genes (Huang da et al. 2009a; Huang da et al. 2009b). Sixteen clusters were observed among the downregulated genes with 42 membrane-associated proteins constituting the largest group. Nine clusters were observed among upregulated genes, which included the keywords Membrane, Cytoskeleton, Microtubule, Metalloprotease, Rap1 signaling and Cell junctions. All of the terms identified among upregulated and downregulated genes are relevant to the regulation of vascular integrity (Dudek and Garcia 2001; Chrzanowska-Wodnicka 2017). Therefore, we analyzed the expression of cell junction molecules in *Gata2^LECKO^* and *miR-126^−/−^* embryos. Claudin5 expression was dramatically downregulated in the lymphatic vessels of E16.5 *miR-126^−/−^* embryos (**Figure 10A, A’, B, B’’**). Additionally, whereas VE-Cadherin was uniformly expressed along the cell junctions of control embryos, it displayed discontinuous expression in *miR-126^−/−^* embryos (**Figure 10A’’, A’’’, B’’, B’’’**). We identified identical defects in VE-Cadherin and Claudin5 expression in E16.5 *Gata2^LECKO^* embryos (**Figure 10C, D**). In addition, VE-Cadherin expression was disorganized in the mesenteric lymphatic vessels of E18.5 *Prox1-CreERT2; Gata2^f/f^* embryos that were exposed to tamoxifen at E14.5 (**Figure 10E, F**).

**Figure 10:**
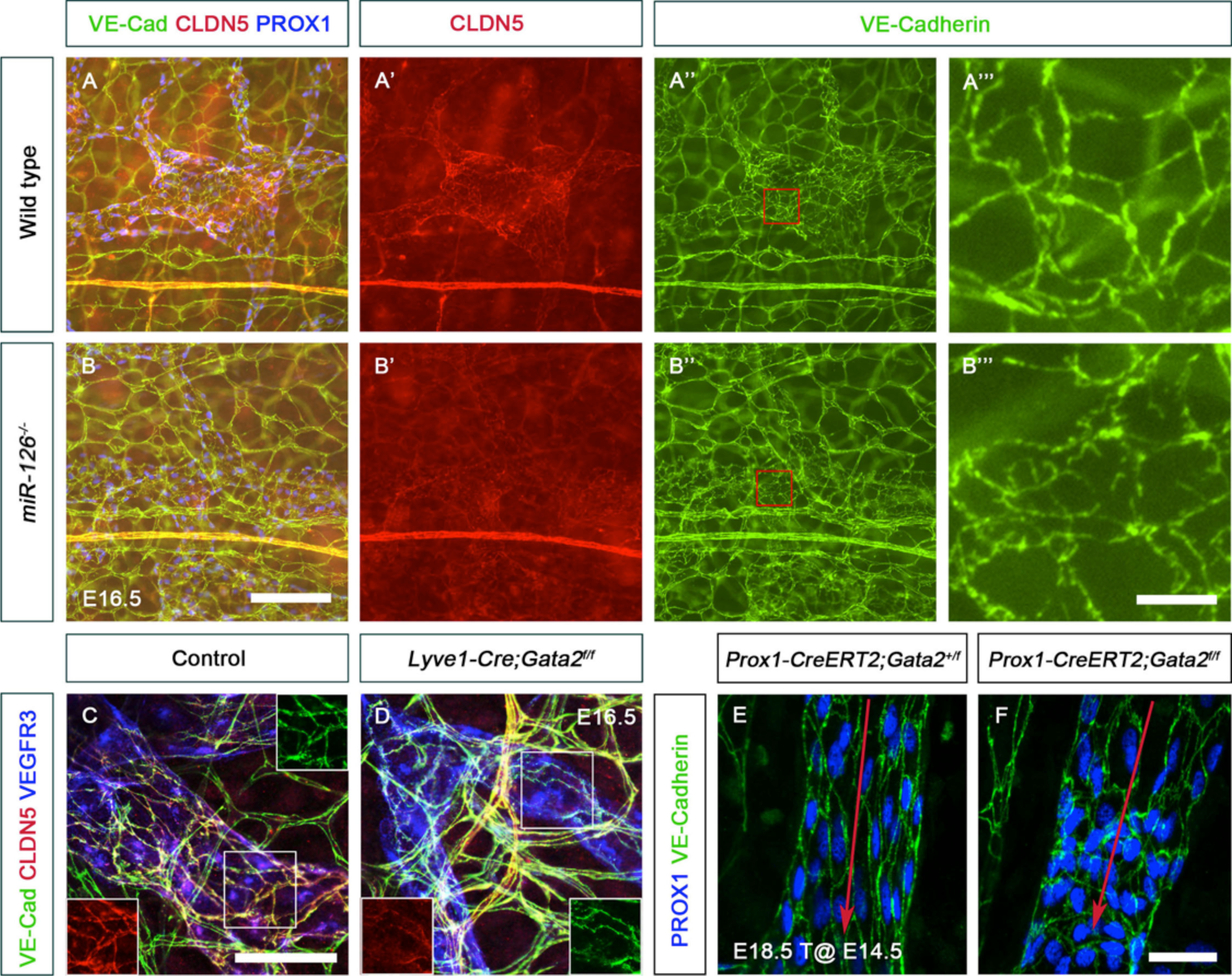
GATA2 and miR-126 are necessary for the expression of cell junction molecules Claudin5 and VE-Cadherin. (A, B) Expression of tight junction molecule Claudin5 (CLDN5) was dramatically downregulated in the dorsal skin lymphatic vessels of E16.5 *miR-126^−/−^* embryos (B’). The intensity of VE-Cadherin staining appeared to be not different between control (A’’) and mutant (B’’) embryos. However, closer inspection revealed that VE-Cadherin was uniformly expressed along the cell boundaries of control embryos (A’’’), but was localized in a “zig zag” pattern in mutant (B’’’) embryos. (C, D) Claudin5 was downregulated (D, left inset) and VE-Cadherin had a defective localization with gaps (D, right inset) in the dorsal skin lymphatic vessels of E16.5 *Lyve1-Cre;Gata2^f/f^* embryos. (E) The LECs of mesenteric lymphatic vessels were elongated in the direction of lymph flow (red arrow) in E18.5 control embryos. VE-Cadherin was uniformly expressed around the cell boundaries of control LECs. In contrast, the LECs were misaligned and VE-Cadherin appeared to be mislocalized in embryos lacking GATA2 (F). Measuring bar: (A, B) 200 µm; (A”’-B’”) 25 µm; (C-F) 50 µm. Statistics: n= 3 embryos per genotype.

Thin sections of LVV-ECs from E12.0 control and *Gata2^LECKO^* embryos did not reveal any obvious differences in VE-Cadherin or Claudin5 expression (**Figure 2- Figure Supplement 1M-P**). However, LVV-ECs delaminate from the walls of veins at E12.0 before reassembling in multiple layers to form mature LVVs at E12.5 (Geng et al. 2016). Such a rapid morphogenesis of LVV-ECs is likely to involve dramatic reorganization of cell junctions. And, the deletion of VE-Cadherin from the lymphatic vasculature was recently reported to prevent the formation of LVs (Hagerling et al. 2018). Hence, we are tempted to speculate that a defect in the reorganization of cell junctions might be the cause of LVV-EC disappearance in E12.5 *Gata2^LECKO^* embryos.

Given that the lymphatic vessels of E16.5 *Gata2^LECKO^* and *miR-126^−/−^* embryos had defective cell junctions, we examined Claudin5 and VE-Cadherin expression in HLECs. Claudin5 and VE-Cadherin were uniformly expressed around the entire periphery in ∼80% of control HLECs (**Figure 11A, B**). In contrast, the intensity of Claudin5 expression was dramatically reduced in GATA2^ΔHLEC^ cells (**Figure 11C**). In addition, the localization of VE-Cadherin was defective in GATA2^ΔHLEC^ cells with numerous gaps (**Figure 11D**, arrowheads). To determine whether miR-126 also influences Claudin5 expression in HLECs, we overexpressed the “miR-126 sponge” (Lechman et al. 2012) in HLECs, and observed a significant reduction in Claudin5 expression and defective VE-Cadherin localization (**Figure 11E-H**). Thus, both GATA2 and *miR-126* are regulators of Claudin5 expression and VE-Cadherin localization *in vitro* and *in vivo*.

**Figure 11:**
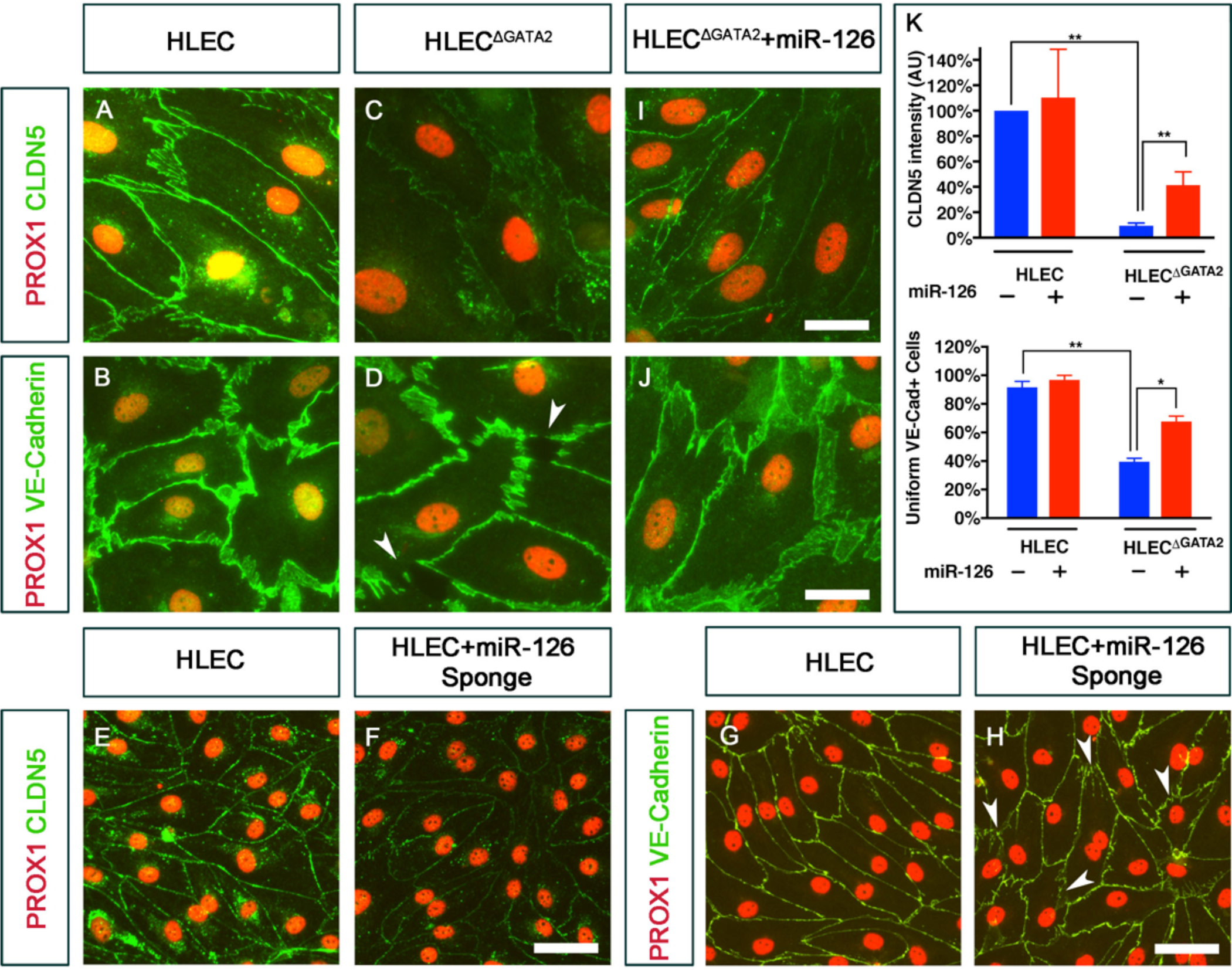
GATA2 regulates the expression of Claudin5 and VE-Cadherin in HLECs in a miR-126 dependent manner. (A) Claudin5 and (B) VE-Cadherin were uniformly expressed in the cell boundaries of control HLECs. In contrast, (C) Claudin5 was dramatically downregulated and (D) VE-Cadherin was expressed in a zig zag pattern (arrowheads) on the cell boundaries of HLEC^ΔGATA2^. (E-H) Expression of a miR-126 sponge using lentivirus strikingly downregulated Claudin5 (F) and caused disruptions in VE-Cadherin localization (H, arrowheads) in HLECs. (I, J) Overexpression of miR-126 using lentiviral particles significantly rescued the expression of Claudin5 (I) and VE-Cadherin (J) expressions in HLEC^ΔGATA2^. (K) The intensity of Claudin5 expression and the number of cells with uniform expression of VE-Cadherin were quantified and graphically shown in K. Measuring bar: (A-F) 25 µm; (H-K) 50 µm. Statistics: (A-D, E-H, I) n = 3; (J) n=2. (**) p<0.01; (*) p<0.05.

To determine whether GATA2 regulates Claudin5 via miR-126, we overexpressed *miR-126* in GATA2^ΔHLEC^ cells using lentiviral particles (Amendola et al. 2009). We observed a partial, yet significant rescue of Claudin5 expression in GATA2^ΔHLEC^ expressing *miR-126*. In addition, miR-126 significantly rescued the localization of VE-Cadherin at the cell junctions (**Figure 11I-G**). These results suggest that GATA2 regulates adherens and tight junctions in LECs through miR-126.

## DISCUSSION

In this work, we have discovered that although GATA2 is not necessary for LVV-EC differentiation, it is required for their maintenance. Further, GATA2 is important for LVV-ECs and LECs to align appropriately with respect to the direction of fluid flow. GATA2 activates the expression of *miR-126* in LVV-ECs and LECs. The lymphatic vascular defects of mice lacking GATA2 or *miR-126* are strikingly similar, and both GATA2 and *miR-126* are necessary for the expression of cell junction molecules Claudin5 and VE-Cadherin. Importantly, *miR-126* could significantly rescue cell junction defects in HLECs lacking GATA2. Based on our results we propose a model in which GATA2 regulates LVV morphogenesis and lymphatic vascular maturation by maintaining proper cell junctions via *miR-126* (**Figure 12**).

**Figure 12:**
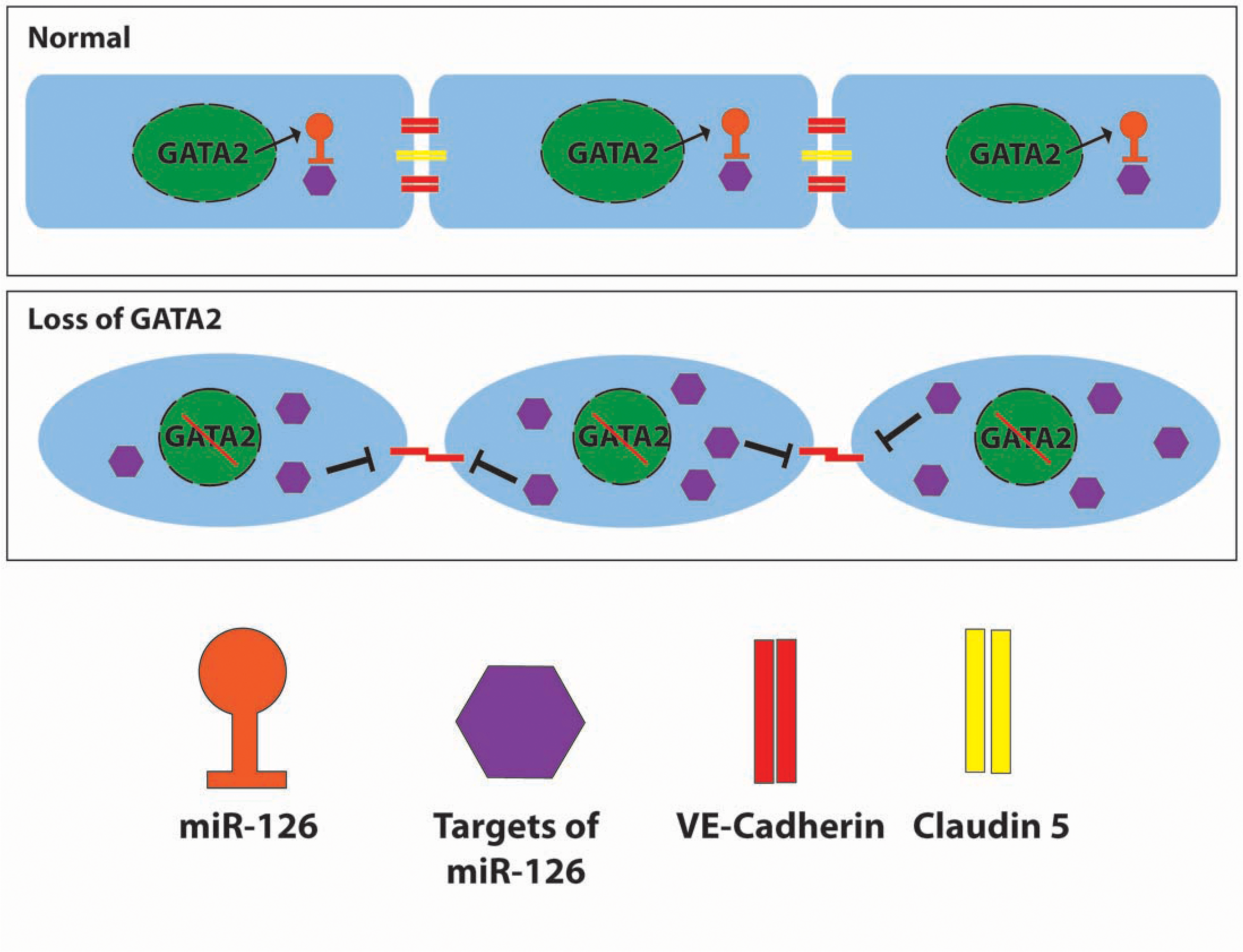
GATA2 regulates endothelial cell junctions through miR-126. Under normal conditions GATA2 enhances the expression of miR-126 in the lymphatic vasculature. The targets of miR-126 are upregulated in the absence of GATA2, which results in the downregulation of Claudin5 and mislocalization of VE-Cadherin. Consequently, lymphatic vascular morphogenesis is defective due to abnormal shape of LECs and LVV-ECs.

Valves normally develop at locations of disturbed flow. Hence, pioneering work by Sabine *et al*. proposed OSS as the most upstream regulator of valve development (Sabine et al. 2012). Significant advances have been made since this initial report. We showed that Wnt/ß-catenin signaling enhances the expression of FOXC2 and GATA2 in response to OSS (Cha et al. 2016), and that PROX1 is necessary for the activity of Wnt/ß-catenin signaling (Cha et al. 2018). Elegant studies have shown that GATA2 enhances the expression of FOXC2 in an OSS-dependent manner (Kazenwadel et al. 2015; Sweet et al. 2015). Despite this knowledge, the precise role of OSS in valve development is unknown. Cytoskeleton, cell-cell and cell-matrix interactions are critical regulators of mechanotransduction (Ingber 2006; Hahn and Schwartz 2009). Therefore, our finding that GATA2 regulates VE-Cadherin and Claudin5 expression through miR-126 provides a mechanistic explanation for GATA2-mediated mechanotransduction. Surprisingly, GATA2 is not necessary for the upregulation of FOXC2 expression in LVV-ECs *in vivo*. Therefore, we suggest that the OSS/GATA2/FOXC2 axis is not necessary for the differentiation of valvular endothelial cells. Instead, OSS might be important for lymphatic vessel patterning and the maintenance of FOXC2 expression in valvular endothelial cells. In line with this model, the mechanosensory ion channel PIEZO1 was recently shown to be necessary for LV development (Nonomura et al. 2018; Choi et al. 2019). Just like GATA2, PIEZO1 is not necessary for the differentiation of LV-ECs (Nonomura et al. 2018). Instead, PIEZO1 maintains LV-ECs and regulates their morphogenesis into LVs.

Several interesting questions arise from our work for future exploration. Recently, Kontarakis et al reported that they do not observe any obvious edema or lymphatic defects in *miR-126^−/−^* embryos (Kontarakis et al. 2018). However, we consistently observe severe edema and lymphatic vascular defects in *miR-126^−/−^* embryos (**Figure 9B)**. The reasons for these phenotypic differences remain to be determined. Neither GATA2 nor *miR-126* regulates Claudin5 at the mRNA level (**Supplementary File 1** and data not shown). Hence, how *miR-126* regulates Claudin5 is currently unknown. The mechanisms behind the abnormal patterning of VE-Cadherin are also not known. It will be of interest to generate mice lacking cell junction molecules VE-Cadherin and Claudin5, to determine if they recapitulate any of the phenotypes of *Gata2^LECKO^* and *miR-126^−/−^* embryos. Future work should also address whether modulators of cell-ECM interaction and cadherin stability such as ADAM19 and MMP15, which are significantly upregulated in shGATA2- and miR-126 sponge-treated HLECs, play physiologically important roles in the development of the lymphatic vasculature.

How GATA2 regulates blood-lymph separation is not yet clear. LVV-ECs do not form until E12.0 (Srinivasan and Oliver 2011; Geng et al. 2016). However, the lymph sacs of E11.5 *Gata2^LECKO^* embryos are blood filled (data not shown). Furthermore, as mentioned previously, *Prox1-CreERT2;Gata2^f/f^* embryos did not have blood-filled lymphatics phenotype despite the absence of LVVs (**Figure 3E, F**). These observations suggest that GATA2 is regulating blood-lymph separation through an LVV-independent mechanism.

Platelet-expressed CLEC2 and LEC-expressed podoplanin play a critical role in blood-lymph separation (Fu et al. 2008; Bertozzi et al. 2010). Expression of podoplanin is not affected in *Gata2^LECKO^* embryos, and platelets are present in *Gata2^LECKO^* embryos (data not shown). Together these results suggest that GATA2 regulates blood-lymph separation independent of the platelet/LEC interaction. As GATA2 regulates the expression of numerous genes in blood endothelial cells, and as LECs originate predominantly from embryonic veins, we are tempted to speculate that GATA2 might be regulating blood-lymphatic separation by maintaining blood vascular integrity during the migration of LECs from the veins.

Only some patients with mutations in GATA2 will develop lymphedema. A subset of mice lacking one allele *Gata2* in endothelial cells display a dilated thoracic duct with reduced lymph flow (Kazenwadel et al. 2015), but they have a normal lifespan and do not develop any obvious symptoms of the human disease such as leukemia, bacterial infections or warts (our unpublished observations). Compared to humans, mice experience less severe gravitational load in the lymphatic vessels of their limbs (Castorena-Gonzalez et al. 2018), which might underlie the absence of obvious lymphatic defects in *Gata2^+/-^* mice. Alternative explanations are also possible. Secondary mutations in *ASXL1* are frequently observed in Emberger syndrome patients who develop MDS/AML (Crispino and Horwitz 2017). Similarly, a “second hit” in the targets of GATA2, such as miR-126, might be required to trigger the onset of lymphedema in human patients. Non-coding RNAs are powerful biomarkers of human disease due to their ability to be detected in bodily fluids (Van Roosbroeck et al. 2013). Whether circulating miR-126 levels might predict the onset of lymphedema in Emberger syndrome patients needs to be determined.

Finally, several miRNA mimics and miRNA inhibitors have entered Phase I, Phase II and preclinical trials and approaches to deliver them are rapidly improving (Rupaimoole and Slack 2017). Hence, we are excited about the possibility that miR-126 might one day be used as a “drug” to treat lymphedema in Emberger syndrome patients.

## MATERIALS AND METHODS

### Antibodies

Primary antibodies for immunohistochemistry: rabbit anti-PROX1 (Cat# 11-002, Angiobio, San Diego, CA, USA), goat anti-human PROX1 (Cat# AF2727, R&D Systems, Minneapolis, MN, USA), sheep anti-mouse FOXC2 (Cat# AF6989, R&D Systems, Minneapolis, MN, USA), goat anti-mouse VEGRF3 (Cat# AF743, R&D Systems, Minneapolis, MN, USA), rat anti-mouse CD31 (Cat# 553370, BD Pharmingen, San Jose, CA, USA), goat anti-mouse ITGA9 (Cat # AF3827, R&D Systems, Minneapolis, MN, USA), rat anti-mouse VE-Cadherin (Cat# 550548, BD Pharmingen, San Jose, CA, USA), hamster anti-mouse PDPN (Cat# 127401, Biolegend, San Diego, CA, USA), rat anti-mouse ITGA5 (Cat #553319, BD Pharmingen, San Jose, CA, USA), goat anti-mouse GATA2 (Cat #AF2046, R&D Systems, Minneapolis, MN, USA), rabbit anti-mouse CX37 (Cat #40-4200, Life technologies, Grand Island, NY, USA), rabbit anti-mouse LAMA5 (Abcam, Cambridge, MA, USA), rabbit anti-human Fibronectin (Cat# ab2413, Abcam, Cambridge, MA, USA), Goat anti-human ANGPT2 (Cat# AF623, R&D Systems, Minneapolis, MN, USA), goat anti-mouse EGFL7 (Cat # AF3089, R&D Systems, Minneapolis, MN, USA), rabbit anti-mouse CLDN5 (Cat #34-1600, Thermo Fisher Scientific, Rockford, IL, USA), rabbit anti-mouse LYVE-1 (Cat # 11-034, Angiobio, San Diego, CA, USA).

Secondary antibodies for immunohistochemistry: Cy3-conjugated donkey anti-rabbit, Cy3-conjugated donkey anti-sheep, and Cy5-conjugated donkey anti-rat antibodies were purchased from Jackson ImmunoResearch Laboratories (West Grove, PA, USA). Alexa 488-conjugated donkey anti-goat, Alexa 488-conjugated goat anti-chicken, and Alexa 488-conjugated donkey anti-rat were purchased from Life Technologies (Grand Island, NY, USA).

Primary antibodies for Western blotting: mouse anti-β-Actin (Cat# A5441, Sigma-Aldrich, St. Louis, MO, USA), goat anti-human PROX1 (Cat# AF2727, R&D Systems, Minneapolis, MN, USA), goat anti-mouse GATA2 (Cat #AF2046, R&D Systems, Minneapolis, MN, USA), goat anti-mouse EGFL7 (Cat # AF3089, R&D Systems, Minneapolis, MN, USA) and mouse anti-human ANGPT2 (Cat# DY623, R&D Systems, Minneapolis, MN, USA), Rabbit anti-human GAPDH (Cat# PAB13195, Abnova, Taipei City, Taiwan, R.O.C).

HRP-conjugated secondary antibodies for Western blotting: goat anti-mouse IgG, goat anti-rabbit IgG, donkey anti-goat IgG, and donkey anti-sheep IgG from Santa Cruz Biotechnology.

### Cells

We used de-identified primary human lymphatic endothelial cells (HLECs) for experiments. HLEC-1 were from Lonza (CC-2812) and were used for RNA-seq analysis with shGATA2. HLEC-2 (Lonza, CC-2516) were used for all other experiments. HLECs were grown on firbonectin-coated plates or glass slide and were maintained in EBM2 media from Lonza. All experiments were conducted using passage 5-6 cells. HLECs were treated as potential biohazards and were handled according to institutional biosafety regulations.

### Chromatin Immunoprecipitation

ChIP assays were performed using EZ-ChIP kit (MilliporeSigma, Burlington, MA, USA) according to the manufacturer’s instructions. Around 1.0 × 10^7^ HLECs were used per ChIP. Briefly, HLECs were grown on culture dish at around 100% confluence. Subsequently, HLECs were fixed in 1% formaldehyde for 10 min at room temperature and glycine at a final concentration of 0.125 M was added for 5 min. Cells were washed with 20 mL of ice cold PBS twice (ten minutes each) and harvested. Cells were lysed and sonicated as described previously described (Cha et al. 2016; Cha et al. 2018).

Chromatin immunoprecipitation was performed using 3.0 µg of goat anti-mouse GATA2 (R&D Systems, Minneapolis, MN, USA) or 1.0 µg of normal goat IgG antibody (R&D Systems, Minneapolis, MN, USA). Q-PCR following ChIP was performed using primers flanking the predicted GATA2 binding sites or control sites of miR-126 promoter.

### Immunohistochemistry of tissues

Immunohistochemistry on sections was done according to our previously published protocols (Cha et al. 2016; Geng et al. 2016; Cha et al. 2018). Briefly, freshly collected embryos were washed in 1X PBS and fixed in 4% paraformaldehyde (PFA) overnight at 4°C. Subsequently, the embryos were washed 3X (10 minutes each) in cold PBS, incubated in 15% sucrose overnight at 4°C and then in 30% sucrose at 4°C until fully submerged in the solution. Embryos were then cryo embedded in OCT solution (Sakura, Tokyo, Japan). 12 µm thick cryosections were prepared using a cryotome (Thermo Fisher Scientific, Model: HM525 NX) and immunohistochemistry was performed using the indicated antibodies. E11.5 embryos were sectioned in a transverse orientation and E12.0-E16.5 embryos were sectioned frontally. Several consecutive sections were analyzed to determine the presence or absence of LVVs and VVs.

Whole mount immunohistochemistry using embryonic skin or guts was performed according to our previous protocol (Cha et al. 2016; Cha et al. 2018). Either whole embryos or isolated guts were washed in 1X PBS and fixed in 1% PFA for 1 hr to overnight (depending on the antibody) at 4°C. Subsequently, the dorsal skins were isolated, washed and samples were immunostained using the iDISCO protocol (Renier et al. 2014). Samples were visualized and analyzed as described previously (Cha et al. 2016; Cha et al. 2018).

### Immunostaining of cells

Cells were fixed in 1% PFA at room temperature for 30 min. Cells were subsequently permeabilized with 0.3 % triton X-100 for 10 min at room temperature, then washed with PBST (PBS + 0.1% Triton-X100) and blocked in 0.5% BSA PBST for 1hour at room temperature. Samples were incubated with primary antibodies at 4°C overnight. Samples were then washed with PBST and incubated with secondary antibodies for 2 hours at room temperature, and then washed with PBST three times (10 min each), mounted and visualized as previously described (Cha et al. 2016; Cha et al. 2018).

### In situ hybridization

We used a kit to detect mmu-miR-126-3p by in situ hybridization (catalog number 339111, Qiagen, Germantown, MD, USA). Briefly, we fixed the embryos in 4% PFA overnight at 4°C. They were then soaked in sucrose, embedded in OCT and sectioned as described above. The sections were fixed in 4% PFA for 10 min at RT and washed in PBS. Subsequent steps were performed according to manufacturer’s instructions.

### Knockdown of GATA2

shGATA2 (TTAACAGGCCACTGACCATGAAGAAGGAA) was cloned into a pLV plasmid. Cyagen Bioscience (Santa Clara, CA, USA) generated the lentiviral particles using LentiPAC 293 cells. HLECs were seeded at 50-60% confluence on fribronectin-coated plates. The following day cells are infected with equal amounts of shcontrol or shGATA2 virus according to manufacturer protocol for 4-6 hrs in Opti-MEM medium and then changed to regular EBM2 media. After 2-3 days cells were harvested with Trizol (Invitrogen, Carlsbad, CA, USA) for RNA-seq study.

### Knockout of GATA2 using CRISPR/Cas9

sgRNA1 (GGTCTGGGTGCAGACGGCAA), sgRNA2 (ATGCCAACCCCGCTCACGCG) and Cas9 were cloned into a pLV plasmid with puromycin selection marker. The translational start site ATG of *GATA2* is located between the recognition sites of sgRNA1 and sgRNA2. Cyagen Bioscience (Santa Clara, CA, USA) generated the lentiviral particles using LentiPAC 293 cells. HLEC-2 were seeded at 50-60% confluence on fibronectin-coated plates. The following day cells were infected with equal amount of control or GATA2 CRIPSPR/cas9 recombinant lentiviral particles according to manufacturer protocol for 4-6 hrs in Opti-MEM medium and then changed to regular EBM2 media. After 24 hrs cells were treated with 0.5µg/mL puromycin to select the cells. After 3 days 0.5µg/mL puromycin treatment almost all non-infected HLECs were dead. We used 5 days of 0.5µg/mL puromycin treatment for selecting GATA2^ΔHLEC^.

A gene-specific primer pair was used that could cover both sgRNA1 and sgRNA2 sequences, generating 330 bp long amplicons. The resulting PCR amplicons were purified using MinElute PCR purification kit (Qiagen). Sequencing library was constructed from 100 ng DNA and approximately 50-100K 300-base read pairs were generated on an Illumina MiSeq platform. GeneWiz Inc performed library preparation, sequencing and bioinformatics analysis.

A total of 66,473 reads were aligned to reference sequence. Sequences that occurred with the frequency of 5 or more were used for further analysis, and a total 63,526 sequences fit this criterion. Indels were detected in 62,864 sequence reads (∼99%). There are 62,808 sequences with deletions. 42,334 reads (67%) harbor a 231 bp deletion between the two targets. 232 bp and 248 bp long deletions were also detected with lower frequencies (13.2%, and 1.2% respectively). There were 2399 sequences (∼3.8%) with two deletions (12 bp and 19 bp) within the sgRNA1, and sgRNA2 respectively.

### Mice

*Prox1^+/Cre^* (Srinivasan et al. 2010)*, Gata2^f/f^* (Charles et al. 2006), *miR-126^−/−^* (Wang et al. 2008)*, Prox1-CreERT2* (Srinivasan et al. 2007), Tg(Prox1-tdTomato) (Gong et al. 2003)*, Lyve1-Cre* (Pham et al. 2010), Tie2-Cre (Kisanuki et al. 2001), *R26^+/tdTomato^* (Madisen et al. 2010) mice were described previously*. Prox1^+/Cre^* mice were maintained in NMRI background. Other mice were maintained in C57BL6 or C57BL6/NMRI mixed backgrounds. We used both male and female mice for the experiments. All mice were housed and handled according to the institutional IACUC protocols.

### miR-126 sponge and miR-126 over expression

pSFFV plasmids to sequester miR-126 or over express miR-126 were reported previously (Amendola et al. 2009; Gentner et al. 2009). Cyagen Bioscience (Santa Clara, CA, USA) generated the lentiviral particles using LentiPAC 293 cells. HLECs were seeded at 50-60% confluence on fribronectin-coated plates or glass slide. The following day cells are infected with equal amounts of control, miR-126 sponge or miR-126 over expression virus for 4-6 hrs according to manufacturer protocol using EBM2 medium and then changed to fresh medium. After 2-3 days cells were harvested for appropriate study.

### miR-126 isolation and quantitative real-time PCR

Micro RNA along with total RNA was isolated from HLECs using QIAzol lysis reagent (Qiagen, Germantown, MD, USA) according to manufacturers instructions. The cDNA was synthesized from total RNA (0.1 – 1.0 µg) with miScript II RT Kit (Qiagen, Germantown, MD, USA). qRT-PCR was performed using miScript SYBR Green PCR Kit (Qiagen, Germantown, MD, USA) in a CFX96 Real-Time System (Bio-Rad, Hercules, CA, USA). miR-126 Expression levels were normalized to *U6.* Predesigned primers for miR-126 and U6 were purchased from Qiagen (cat # 218300).

### RNA-seq analysis

Total RNA was purified from HLECs infected with shGATA2 or control shRNA expressing lentivirus particles. RNA was subjected to ribosomal RNA depletion followed by Truseq stranded total RNA library preparation according to the manufacturer’s instruction (Illumina). RNA from miR-126-sponge treated HLEC-2 were processed using NEB Ultra II directional RNA Library kit for Illumina. The resulting RNA-seq libraries were analyzed on the Illumina HiSeq sequencing platform. The obtained sequencing reads were mapped with the bowtie2 algorithm using the RefSeq annotations (hg19 genome build) (Langmead and Salzberg 2012). We utilized the RNA-seq analysis work flow within the Partek Genomics Suite (Partek Incorporated) for quantitation and statistical analysis (ANOVA) of the transcriptome data. We identified those transcripts that exhibited statistically significant differential expression in the shGata2 samples compared to the shControl samples. We rank ordered the two lists based on the expression level and magnitude of change. Using these rank-ordered list, we performed gene ontology (GO) analysis for enriched biological terms (Eden et al. 2009). The genes commonly regulated by GATA2 and miR-126 were analyzed using the functional annotation platform of DAVID (Huang da et al. 2009a; Huang da et al. 2009b).

### Scanning Electron Microscopy

SEM was performed according to our previous protocol (Geng et al. 2016; Geng and Srinivasan 2018).

### Statistical Analysis

For biochemical analysis n indicates the number of times the experiments were performed and for histological analysis n indicate the number of embryos analyzed per genotype. VE-Cadherin expression analysis in GATA2^ΔHLEC^ following miR-126 overexpression was performed twice. All other experiments were performed at least three times or more. Data were presented as mean ± SEM. GraphPad Prism 7 software was used to perform the statistical analysis. Data were analyzed by the unpaired, two-tailed, student’s *t* test. *P* value < 0.05 was considered significant.

### Western blot

Control HLEC or GATA2^ΔHLEC^ were grown in 12-well plates at ∼ 100% confluency. Cells were harvested with lysis buffer and Western blots were performed using standard protocol.

## ACKNOWLEDGEMENT

This work is supported by NIH/NHLBI (R01HL131652 to RSS and THK; R01HL133216 to RSS and HC), NIH/NIGMS COBRE (P20 GM103441 to XG; PI: Dr. McEver), Oklahoma Center for Adult Stem Cell Research (4340) to RSS and American Heart Association (16PRE31190025 for MRM). We thank Dr. Angela Andersen (Life Science Editors) for editorial assistance, Drs. Eric Lehmann and John Dick (University health Network, Toronto, Canada), and Luigi Naldini (San Raffaele Telethon Institute for Gene Therapy, Milan, Italy) for miR-126 lentiviral constructs, and Mrs. Lisa Whitworth and Mr. Brent Johnson (Oklahoma State University) for SEM.

## AUTHOR CONTRIBUTIONS

MRM, XG, YCH, BC, LC, YK, GM, THK, K-CL and RSS performed experiments; DC, HC, Y-KH, SC and JDE provided critical reagents; MRM, XG, and RSS designed the experiments and wrote the manuscript; all authors provided input in designing the experiments and in writing the manuscript.

---

**Figure 2- Figure Supplement 1:**
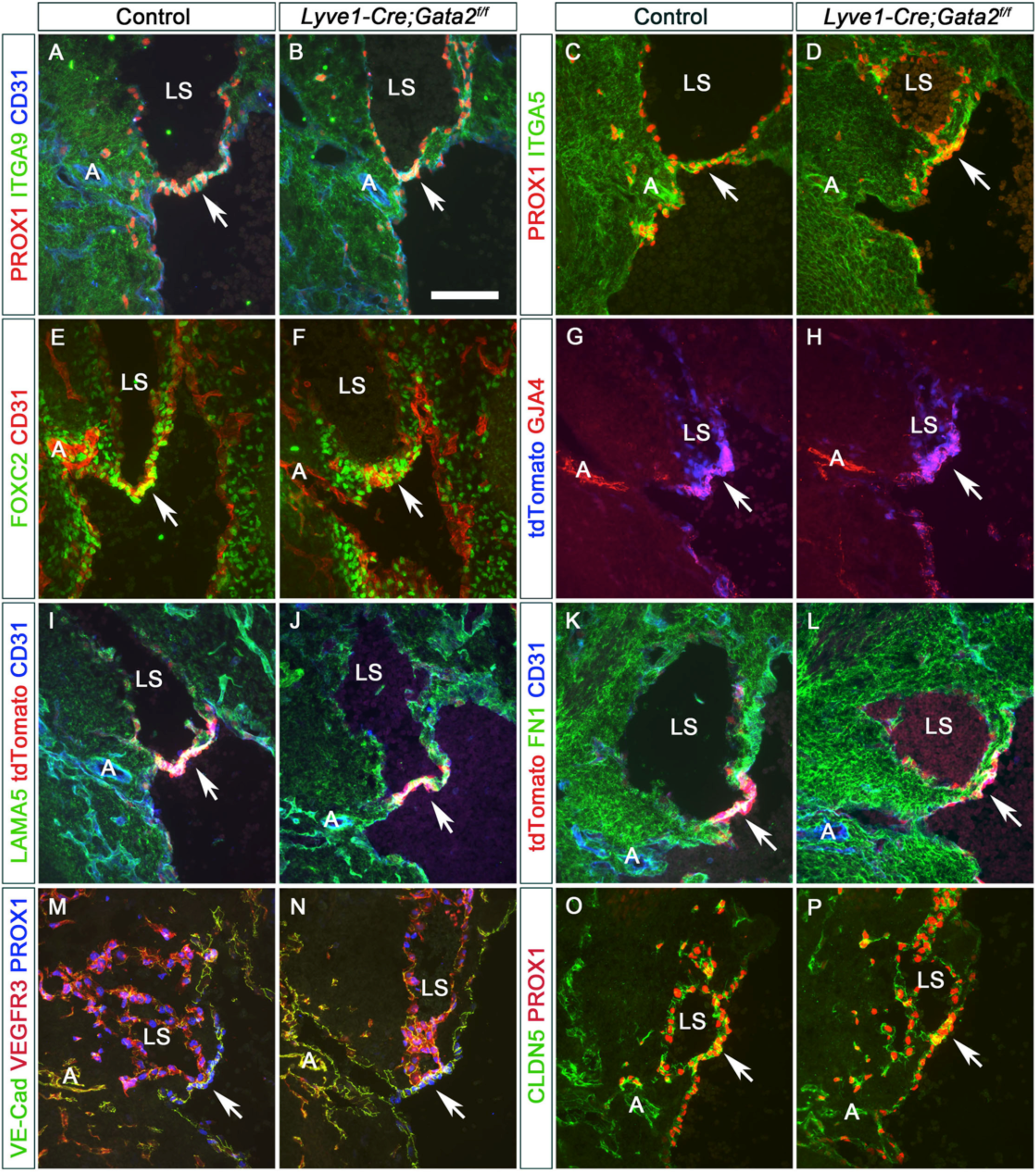
LVV-ECs, ECM and cell junction molecules are normally expressed in E12.0 embryos lacking GATA2. E12.0 Wild type and *Lyve1-Cre;Gata2^f/f^* littermates or Tg(Prox1-tdTomato) and Tg(Prox1-tdTomato); *Lyve1-Cre;Gata2^f/f^* littermates were frontally sectioned and IHC was performed for LVV-EC markers (PROX1, tdTomato, FOXC2, ITGA5, ITGA9), ECM components (FN1, LAMA5), and cell junction proteins (CD31, VE-Cadherin, GJA4, CLDN5). No obvious differences were observed between control and mutant samples. Abbreviations: A, artery; LS, lymph sac. Measuring bar: (A-P) 100 µm. Statistics: n=3 embryos and 6 LVV complexes per genotype per antibody.

**Figure 2- Figure Supplement 2:**
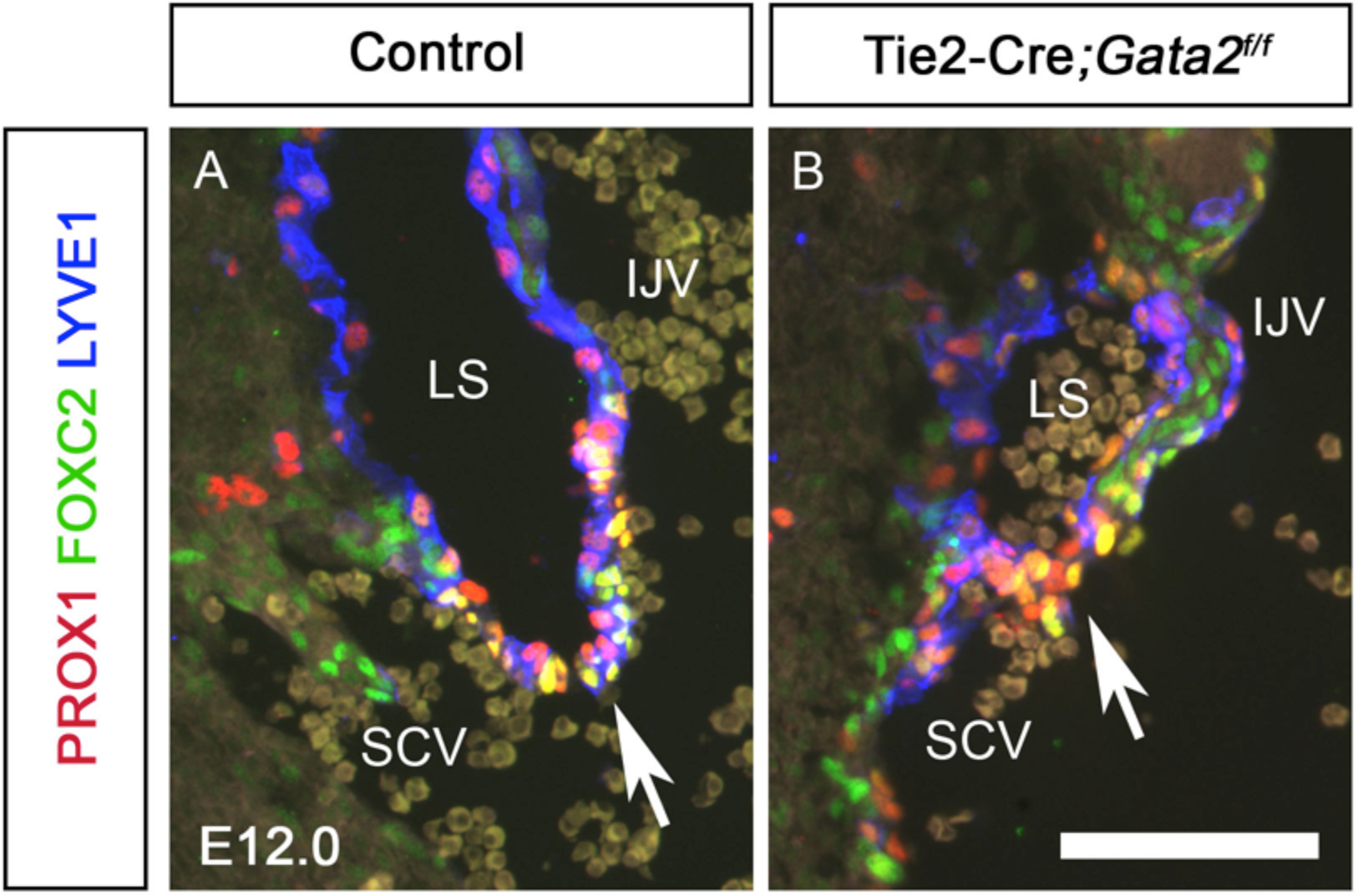
LVV-ECs are present in E12.0 Tie2-Cre;*Gata2^f/f^* embryos. PROX1^+^FOXC2^+^ LVV-ECs are present in both E12.0 control (A, arrow) and mutant embryos lacking GATA2 in all endothelial cells (B, arrow). Abbreviations: IJV, internal jugular vein; SCV, subclavian vein; LS, lymph sac. Measuring bar: (A, B) 100 µm. Statistics: n=3 embryos and 6 LVV complexes per genotype.

**Figure 2- Figure Supplement 3:**
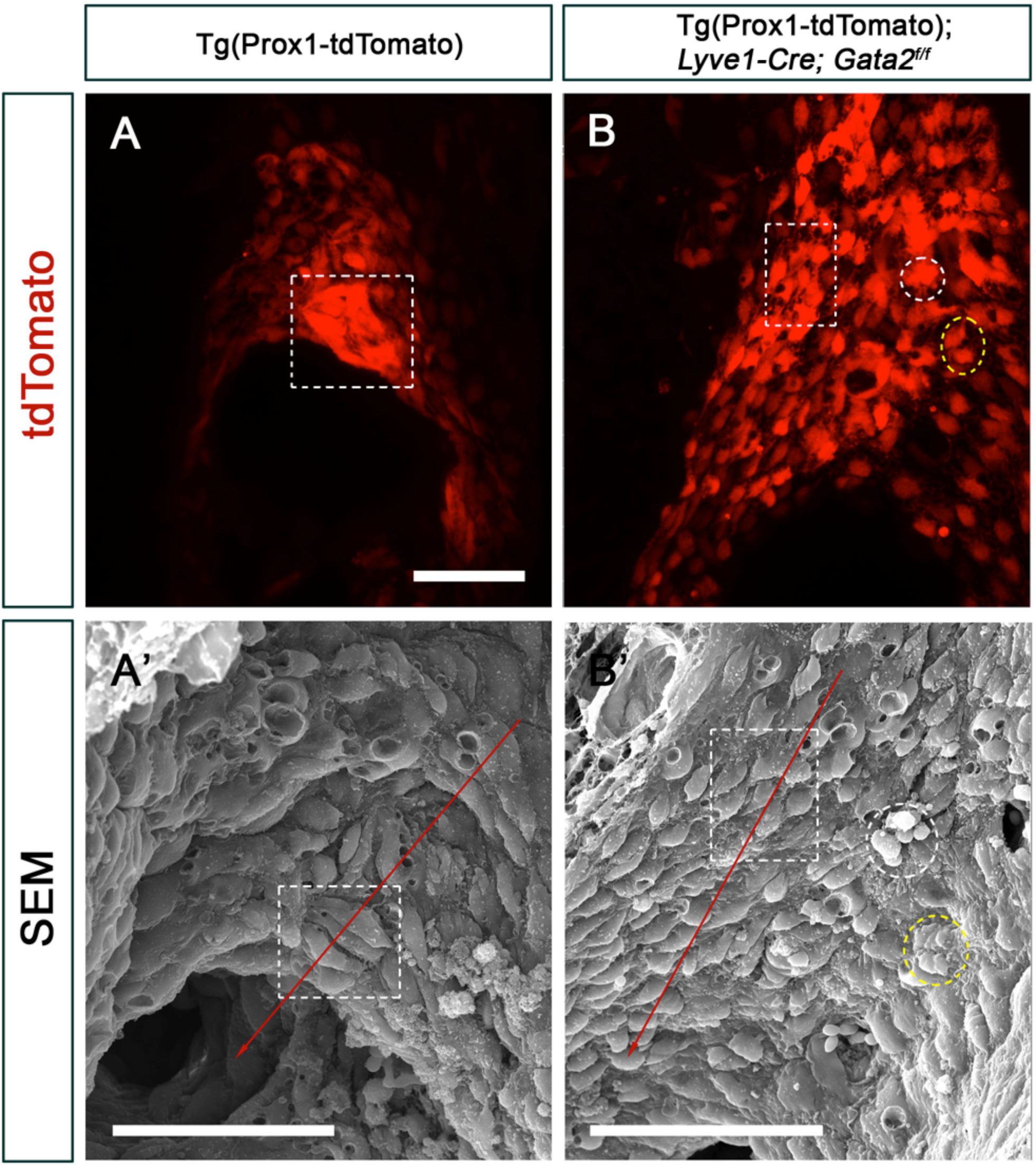
LVV-ECs of E12.0 *Gata2^LECKO^* embryos are dysplastic and do not align properly with respect to blood flow. (A, B) E12.0 Tg(Prox1-tdTomato) or Tg(Prox1-tdTomato); *Lyve1-Cre; Gata2^f/f^* embryos were sagittally sectioned along the internal jugular vein and the tdTomato^+^ LVV-ECs were imaged by confocal microscopy. The same samples were then reprocessed and analyzed by SEM (A’, B’). The cells in fluorescent microscopy images that correlate with cells in the SEM images are outlined. (A’) LVV-ECs of control embryos were elongated and compactly aggregated with each other. They were also aligned perpendicular to the direction of blood flow (red arrow). (B’) In contrast, LVV-ECs of mutant embryos appear round and dispersed (see cells within the rectangular box) and appeared to be aligned parallel to the flow. In addition, several dysplastic cells were also seen (within white and yellow circles). Measuring bar: (A-B) 50 µm; (A’) 40 µm; (B’) 50 µm. Statistics: n=3 embryos, 6 LVV complexes per genotype.

**Figure 5- Figure Supplement 1:**
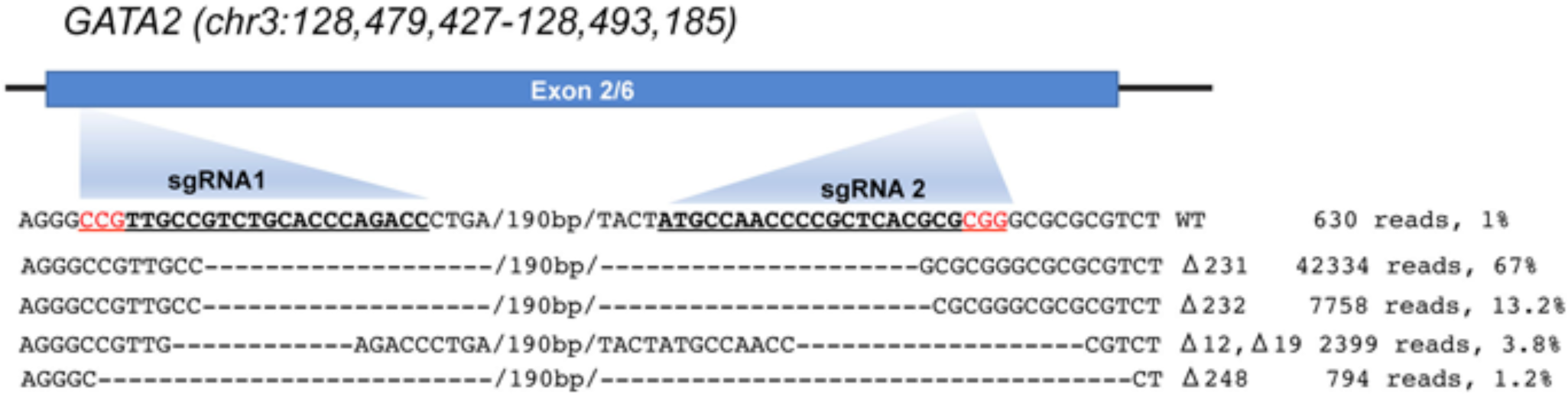
Efficient deletion of GATA2 from HLECs by CRISPR/Cas9. The second exon (out of six total exons) of *GATA2* is schematically shown. Sequences corresponding to the sgRNAs are in bold and are underlined. The PAM sequences are in red. The ATG of GATA2 is located in between sgRNA1 and sgRNA2. HLEC-2 were infected with lentiviruses expressing Cas9 and the two sgRNAs and selected using puromycin. The region between sgRNA1 and sgRNA2 was PCR amplified and was sequenced using NGS approach. The sequence and contribution of major indels is depicted. These results demonstrate that ∼99% of DNA fragments have large deletions in exon 2 of *GATA2*. Statistics: Sequencing was performed using one batch of selected cells (out of 3 independently selected batches).

## REFERENCES

Amendola M, Passerini L, Pucci F, Gentner B, Bacchetta R, Naldini L. 2009. Regulated and multiple miRNA and siRNA delivery into primary cells by a lentiviral platform. Molecular therapy : the journal of the American Society of Gene Therapy 17: 1039–1052.

Bertozzi CC, Schmaier AA, Mericko P, Hess PR, Zou Z, Chen M, Chen CY, Xu B, Lu MM, Zhou D et al. 2010. Platelets regulate lymphatic vascular development through CLEC-2-SLP-76 signaling. Blood 116: 661–670.

Brouillard P, Boon L, Vikkula M. 2014. Genetics of lymphatic anomalies. The Journal of clinical investigation 124: 898–904.

Castorena-Gonzalez JA, Zawieja SD, Li M, Srinivasan RS, Simon AM, de Wit C, de la Torre R, Martinez-Lemus LA, Hennig GW, Davis MJ. 2018. Mechanisms of Connexin-Related Lymphedema. Circulation research 123: 964–985.

Cha B, Geng X, Mahamud MR, Fu J, Mukherjee A, Kim Y, Jho EH, Kim TH, Kahn ML, Xia L et al. 2016. Mechanotransduction activates canonical Wnt/beta-catenin signaling to promote lymphatic vascular patterning and the development of lymphatic and lymphovenous valves. Genes Dev 30: 1454–1469.

Cha B, Geng X, Mahamud MR, Zhang JY, Chen L, Kim W, Jho EH, Kim Y, Choi D, Dixon JB et al. 2018. Complementary Wnt Sources Regulate Lymphatic Vascular Development via PROX1-Dependent Wnt/beta-Catenin Signaling. Cell reports 25: 571–584 e575.

Charles MA, Saunders TL, Wood WM, Owens K, Parlow AF, Camper SA, Ridgway EC, Gordon DF. 2006. Pituitary-specific Gata2 knockout: effects on gonadotrope and thyrotrope function. Molecular endocrinology (Baltimore, Md) 20: 1366–1377.

Charpentier MS, Christine KS, Amin NM, Dorr KM, Kushner EJ, Bautch VL, Taylor JM, Conlon FL. 2013. CASZ1 promotes vascular assembly and morphogenesis through the direct regulation of an EGFL7/RhoA-mediated pathway. Developmental cell 25: 132–143.

Chen H, Griffin C, Xia L, Srinivasan RS. 2014. Molecular and cellular mechanisms of lymphatic vascular maturation. Microvasc Res 96: 16–22.

Choi D, Park E, Jung E, Cha B, Lee S, Yu J, Kim PM, Lee S, Hong YJ, Koh CJ et al. 2019. Piezo1 incorporates mechanical force signals to genetic program that governs lymphatic valve development and maintenance. JCI Insight.

Chrzanowska-Wodnicka M. 2017. Rap1 in endothelial biology. Curr Opin Hematol 24: 248–255.

Craven SE, Lim KC, Ye W, Engel JD, de Sauvage F, Rosenthal A. 2004. Gata2 specifies serotonergic neurons downstream of sonic hedgehog. Development 131: 1165–1173.

Crispino JD, Horwitz MS. 2017. GATA factor mutations in hematologic disease. Blood 129: 2103–2110.

Dellinger M, Hunter R, Bernas M, Gale N, Yancopoulos G, Erickson R, Witte M. 2008. Defective remodeling and maturation of the lymphatic vasculature in Angiopoietin-2 deficient mice. Developmental biology 319: 309–320.

Donadieu J, Lamant M, Fieschi C, de Fontbrune FS, Caye A, Ouachee M, Beaupain B, Bustamante J, Poirel HA, Isidor B et al. 2018. Natural history of GATA2 deficiency in a survey of 79 French and Belgian patients. Haematologica 103: 1278–1287.

Dudek SM, Garcia JG. 2001. Cytoskeletal regulation of pulmonary vascular permeability. J Appl Physiol (1985) 91: 1487–1500.

Eden E, Navon R, Steinfeld I, Lipson D, Yakhini Z. 2009. GOrilla: a tool for discovery and visualization of enriched GO terms in ranked gene lists. BMC Bioinformatics 10: 48.

Emberger JM, Navarro M, Dejean M, Izarn P. 1979. [Deaf-mutism, lymphedema of the lower limbs and hematological abnormalities (acute leukemia, cytopenia) with autosomal dominant transmission]. J Genet Hum 27: 237–245.

Fish JE, Santoro MM, Morton SU, Yu S, Yeh RF, Wythe JD, Ivey KN, Bruneau BG, Stainier DY, Srivastava D. 2008. miR-126 regulates angiogenic signaling and vascular integrity. Developmental cell 15: 272–284.

Frye M, Taddei A, Dierkes C, Martinez-Corral I, Fielden M, Ortsater H, Kazenwadel J, Calado DP, Ostergaard P, Salminen M et al. 2018. Matrix stiffness controls lymphatic vessel formation through regulation of a GATA2-dependent transcriptional program. Nature communications 9: 1511.

Fu J, Gerhardt H, McDaniel JM, Xia B, Liu X, Ivanciu L, Ny A, Hermans K, Silasi-Mansat R, McGee S et al. 2008. Endothelial cell O-glycan deficiency causes blood/lymphatic misconnections and consequent fatty liver disease in mice. The Journal of clinical investigation 118: 3725–3737.

Gale NW, Thurston G, Hackett SF, Renard R, Wang Q, McClain J, Martin C, Witte C, Witte MH, Jackson D et al. 2002. Angiopoietin-2 is required for postnatal angiogenesis and lymphatic patterning, and only the latter role is rescued by Angiopoietin-1. Developmental cell 3: 411–423.

Geng X, Cha B, Mahamud MR, Lim KC, Silasi-Mansat R, Uddin MK, Miura N, Xia L, Simon AM, Engel JD et al. 2016. Multiple mouse models of primary lymphedema exhibit distinct defects in lymphovenous valve development. Developmental biology 409: 218–233.

Geng X, Srinivasan RS. 2018. Correlative Fluorescence and Scanning Electron Microscopy to Study Lymphovenous Valve Development. Methods in molecular biology 1846: 85–96.

Gentner B, Schira G, Giustacchini A, Amendola M, Brown BD, Ponzoni M, Naldini L. 2009. Stable knockdown of microRNA in vivo by lentiviral vectors. Nat Methods 6: 63–66.

Gong S, Zheng C, Doughty ML, Losos K, Didkovsky N, Schambra UB, Nowak NJ, Joyner A, Leblanc G, Hatten ME et al. 2003. A gene expression atlas of the central nervous system based on bacterial artificial chromosomes. Nature 425: 917–925.

Hagerling R, Hoppe E, Dierkes C, Stehling M, Makinen T, Butz S, Vestweber D, Kiefer F. 2018. Distinct roles of VE-cadherin for development and maintenance of specific lymph vessel beds. The EMBO journal 37.

Hahn C, Schwartz MA. 2009. Mechanotransduction in vascular physiology and atherogenesis. Nature reviews Molecular cell biology 10: 53–62.

Hartmann D, Fiedler J, Sonnenschein K, Just A, Pfanne A, Zimmer K, Remke J, Foinquinos A, Butzlaff M, Schimmel K et al. 2016. MicroRNA-Based Therapy of GATA2-Deficient Vascular Disease. Circulation 134: 1973–1990.

Huang da W, Sherman BT, Lempicki RA. 2009a. Bioinformatics enrichment tools: paths toward the comprehensive functional analysis of large gene lists. Nucleic acids research 37: 1–13.

Huang da W, Sherman BT, Lempicki RA. 2009b. Systematic and integrative analysis of large gene lists using DAVID bioinformatics resources. Nature protocols 4: 44–57.

Ingber DE. 2006. Cellular mechanotransduction: putting all the pieces together again. FASEB J 20: 811–827.

Janardhan HP, Milstone ZJ, Shin M, Lawson ND, Keaney JF, Jr., Trivedi CM. 2017. Hdac3 regulates lymphovenous and lymphatic valve formation. The Journal of clinical investigation 127: 4193–4206.

Kazenwadel J, Betterman KL, Chong CE, Stokes PH, Lee YK, Secker GA, Agalarov Y, Demir CS, Lawrence DM, Sutton DL et al. 2015. GATA2 is required for lymphatic vessel valve development and maintenance. The Journal of clinical investigation 125: 2979–2994.

Kazenwadel J, Secker GA, Liu YJ, Rosenfeld JA, Wildin RS, Cuellar-Rodriguez J, Hsu AP, Dyack S, Fernandez CV, Chong CE et al. 2012. Loss-of-function germline GATA2 mutations in patients with MDS/AML or MonoMAC syndrome and primary lymphedema reveal a key role for GATA2 in the lymphatic vasculature. Blood 119: 1283–1291.

Khandekar M, Brandt W, Zhou Y, Dagenais S, Glover TW, Suzuki N, Shimizu R, Yamamoto M, Lim KC, Engel JD. 2007. A Gata2 intronic enhancer confers its pan-endothelia-specific regulation. Development 134: 1703–1712.

Kisanuki YY, Hammer RE, Miyazaki J, Williams SC, Richardson JA, Yanagisawa M. 2001. Tie2-Cre transgenic mice: a new model for endothelial cell-lineage analysis in vivo. Developmental biology 230: 230–242.

Kontarakis Z, Rossi A, Ramas S, Dellinger MT, Stainier DYR. 2018. Mir-126 is a conserved modulator of lymphatic development. Developmental biology.

Kuhnert F, Mancuso MR, Hampton J, Stankunas K, Asano T, Chen CZ, Kuo CJ. 2008. Attribution of vascular phenotypes of the murine Egfl7 locus to the microRNA miR-126. Development 135: 3989–3993.

Lagos-Quintana M, Rauhut R, Yalcin A, Meyer J, Lendeckel W, Tuschl T. 2002. Identification of tissue-specific microRNAs from mouse. Current biology : CB 12: 735–739.

Langmead B, Salzberg SL. 2012. Fast gapped-read alignment with Bowtie 2. Nat Methods 9: 357–359.

Lechman ER, Gentner B, van Galen P, Giustacchini A, Saini M, Boccalatte FE, Hiramatsu H, Restuccia U, Bachi A, Voisin V et al. 2012. Attenuation of miR-126 activity expands HSC in vivo without exhaustion. Cell stem cell 11: 799–811.

Lim KC, Hosoya T, Brandt W, Ku CJ, Hosoya-Ohmura S, Camper SA, Yamamoto M, Engel JD. 2012. Conditional Gata2 inactivation results in HSC loss and lymphatic mispatterning. The Journal of clinical investigation 122: 3705–3717.

Madisen L, Zwingman TA, Sunkin SM, Oh SW, Zariwala HA, Gu H, Ng LL, Palmiter RD, Hawrylycz MJ, Jones AR et al. 2010. A robust and high-throughput Cre reporting and characterization system for the whole mouse brain. Nat Neurosci 13: 133–140.

Mammoto A, Connor KM, Mammoto T, Yung CW, Huh D, Aderman CM, Mostoslavsky G, Smith LE, Ingber DE. 2009. A mechanosensitive transcriptional mechanism that controls angiogenesis. Nature 457: 1103–1108.

Muzumdar MD, Tasic B, Miyamichi K, Li L, Luo L. 2007. A global double-fluorescent Cre reporter mouse. Genesis 45: 593–605.

Nonomura K, Lukacs V, Sweet DT, Goddard LM, Kanie A, Whitwam T, Ranade SS, Fujimori T, Kahn ML, Patapoutian A. 2018. Mechanically activated ion channel PIEZO1 is required for lymphatic valve formation. Proceedings of the National Academy of Sciences of the United States of America.

Ostergaard P, Simpson MA, Connell FC, Steward CG, Brice G, Woollard WJ, Dafou D, Kilo T, Smithson S, Lunt P et al. 2011. Mutations in GATA2 cause primary lymphedema associated with a predisposition to acute myeloid leukemia (Emberger syndrome). Nature genetics 43: 929–931.

Parker LH, Schmidt M, Jin SW, Gray AM, Beis D, Pham T, Frantz G, Palmieri S, Hillan K, Stainier DY et al. 2004. The endothelial-cell-derived secreted factor Egfl7 regulates vascular tube formation. Nature 428: 754–758.

Pham TH, Baluk P, Xu Y, Grigorova I, Bankovich AJ, Pappu R, Coughlin SR, McDonald DM, Schwab SR, Cyster JG. 2010. Lymphatic endothelial cell sphingosine kinase activity is required for lymphocyte egress and lymphatic patterning. The Journal of experimental medicine 207: 17–27.

Renier N, Wu Z, Simon DJ, Yang J, Ariel P, Tessier-Lavigne M. 2014. iDISCO: a simple, rapid method to immunolabel large tissue samples for volume imaging. Cell 159: 896–910.

Rupaimoole R, Slack FJ. 2017. MicroRNA therapeutics: towards a new era for the management of cancer and other diseases. Nature reviews Drug discovery 16: 203–222.

Sabine A, Agalarov Y, Maby-El Hajjami H, Jaquet M, Hagerling R, Pollmann C, Bebber D, Pfenniger A, Miura N, Dormond O et al. 2012. Mechanotransduction, PROX1, and FOXC2 cooperate to control connexin37 and calcineurin during lymphatic-valve formation. Developmental cell 22: 430–445.

Souma T, Thomson BR, Heinen S, Carota IA, Yamaguchi S, Onay T, Liu P, Ghosh AK, Li C, Eremina V et al. 2018. Context-dependent functions of angiopoietin 2 are determined by the endothelial phosphatase VEPTP. Proceedings of the National Academy of Sciences of the United States of America 115: 1298–1303.

Spinner MA, Sanchez LA, Hsu AP, Shaw PA, Zerbe CS, Calvo KR, Arthur DC, Gu W, Gould CM, Brewer CC et al. 2014. GATA2 deficiency: a protean disorder of hematopoiesis, lymphatics, and immunity. Blood 123: 809–821.

Srinivasan RS, Dillard ME, Lagutin OV, Lin FJ, Tsai S, Tsai MJ, Samokhvalov IM, Oliver G. 2007. Lineage tracing demonstrates the venous origin of the mammalian lymphatic vasculature. Genes Dev 21: 2422–2432.

Srinivasan RS, Geng X, Yang Y, Wang Y, Mukatira S, Studer M, Porto MP, Lagutin O, Oliver G. 2010. The nuclear hormone receptor Coup-TFII is required for the initiation and early maintenance of Prox1 expression in lymphatic endothelial cells. Genes Dev 24: 696–707.

Srinivasan RS, Oliver G. 2011. Prox1 dosage controls the number of lymphatic endothelial cell progenitors and the formation of the lymphovenous valves. Genes Dev 25: 2187–2197.

Sweet DT, Jimenez JM, Chang J, Hess PR, Mericko-Ishizuka P, Fu J, Xia L, Davies PF, Kahn ML. 2015. Lymph flow regulates collecting lymphatic vessel maturation in vivo. The Journal of clinical investigation 125: 2995–3007.

Tammela T, Alitalo K. 2010. Lymphangiogenesis: Molecular mechanisms and future promise. Cell 140: 460–476.

Van Roosbroeck K, Pollet J, Calin GA. 2013. miRNAs and long noncoding RNAs as biomarkers in human diseases. Expert review of molecular diagnostics 13: 183–204.

Wang S, Aurora AB, Johnson BA, Qi X, McAnally J, Hill JA, Richardson JA, Bassel-Duby R, Olson EN. 2008. The endothelial-specific microRNA miR-126 governs vascular integrity and angiogenesis. Developmental cell 15: 261–271.

Zhou Q, Anderson C, Hanus J, Zhao F, Ma J, Yoshimura A, Wang S. 2016. Strand and Cell Type-specific Function of microRNA-126 in Angiogenesis. Molecular therapy : the journal of the American Society of Gene Therapy 24: 1823–1835.

Zhou Y, Lim KC, Onodera K, Takahashi S, Ohta J, Minegishi N, Tsai FY, Orkin SH, Yamamoto M, Engel JD. 1998. Rescue of the embryonic lethal hematopoietic defect reveals a critical role for GATA-2 in urogenital development. The EMBO journal 17: 6689–6700.

Zhou Y, Yamamoto M, Engel JD. 2000. GATA2 is required for the generation of V2 interneurons. Development 127: 3829–3838.

